# Bimodular architecture of bacterial effector SAP05 drives ubiquitin-independent targeted protein degradation

**DOI:** 10.1101/2023.06.19.545293

**Authors:** Qun Liu, Abbas Maqbool, Federico G. Mirkin, Yeshveer Singh, Clare E. M. Stevenson, David M. Lawson, Sophien Kamoun, Weijie Huang, Saskia A. Hogenhout

## Abstract

In eukaryotes, targeted protein degradation (TPD) typically depends on a series of interactions among ubiquitin ligases that transfer ubiquitin molecules to substrates leading to degradation by the 26S proteasome. We previously discovered that the bacterial effector protein SAP05 mediates ubiquitin-independent TPD. SAP05 forms a ternary complex via interactions with the von Willebrand Factor Type A (vWA) domain of the proteasomal ubiquitin receptor Rpn10 and the Zinc-finger (ZnF) domains of the SQUAMOSA-PROMOTER BINDING PROTEIN-LIKE (SPL) and GATA BINDING FACTOR (GATA) transcription factors (TFs). This leads to direct TPD of the TFs by the 26S proteasome. Here, we report the crystal structures of the SAP05-vWA complex at 2.17 Å resolution and of the SAP05-ZnF(SPL5) complex at 2.20 Å resolution. Structural analyses revealed that SAP05 displays a unique bimodular architecture with two distinct non-overlapping surfaces, a ‘loop surface’ with three protruding loops that form electrostatic interactions with ZnF, and a ‘sheet surface’ featuring two β-sheets, loops and ⍺-helices that establish polar interactions with vWA. SAP05 binding to ZnF TFs involves single amino acids responsible for multiple contacts, while SAP05 binding to vWA is more stable due to the necessity of multiple mutations to break the interaction. In addition, positioning of the SAP05 complex on the 26S proteasome points to a mechanism of protein degradation. Collectively, our findings demonstrate how a small bacterial bimodular protein can bypass the canonical UPS cellular proteolysis pathway, enabling ubiquitin-independent TPD in eukaryotic cells. This knowledge holds significant potential for the creation of novel TPD technologies.

## Introduction

Pathogenic bacteria secrete effector proteins that manipulate host cell processes, aiding in bacterial survival and often resulting in disease (Hogenhout et al. 2009). These effector proteins are a source of biochemical innovation and include some of the most remarkable proteins known to function inside host cells. Notable examples include Transcription Activator-Like (TAL) effectors, derived from plant pathogenic bacteria, which can be engineered to bind specific DNA sequences (Bogdanove and Voytas 2011), type-III effector proteins from animal parasitic bacteria, capable of re-engineering kinase pathways (Wei et al. 2012), and the CRISPR/Cas systems—immune defenses against bacteriophages, now harnessed as groundbreaking genetic manipulation tools (Mojica and Rodriguez-Valera 2016). In addition, we recently discovered that SAP05 effectors of bacterial phytoplasma pathogens enable the degradation of structured proteins by the 26S proteasome in a ubiquitin-independent manner (Huang et al., 2021). Similar to other bacterial effectors, the utility of SAP05 proteins can extend beyond the realm of natural biology, as they have the potential to serve as tools in biotechnology and biomedicine. However, unraveling the structure-function relationships and underlying mechanisms of SAP05 bacterial effectors is necessary for paving the way for their applications in synthetic biology.

Efficient and selective protein degradation is crucial for all living organisms (Barrett, 2001; Rousseau and Bertolotti, 2018). A significant portion of energy in cells is devoted to the process of protein degradation. This expenditure of energy is vital for cells to adapt and react to their surrounding environment (Bard et al., 2018; Bhattacharyya et al., 2014; Ciechanover, 2017; Collins and Goldberg, 2017; Finley, 2009; Murata et al., 2009). Protein degradation is executed by a diverse family of enzymes, proteases, that hydrolyse peptide bonds (Barrett, 2001). To prevent the destruction of proteins not destined for degradation and avoid the accidental disruption of the cellular proteome, protein degradation must be spatially and temporally controlled (Baumeister et al., 1998; Davis et al., 2021; Inobe and Matouschek, 2014).

The 26S proteasome is such a self-compartmentalizing device. It is a highly sophisticated complex with distinct proteolytic activities able to degrade proteins selectively and efficiently in all eukaryotes (Baumeister et al., 1998; Tanaka, 2009). With 33 different subunits and approximately 2.5 MDa in mass, it is the largest ATP-dependent protease machinery in the cell (Darwin, 2009; Inobe and Matouschek, 2014; Müller and Weber-Ban, 2019). The subunits are organized in two particles: the catalytic 20S core particle (CP) and the 19S regulatory particle (RP). The 20S CP is a cylindrical complex containing four heptameric rings with multiple catalytic β subunits and can degrade intrinsically disordered proteins by itself, including proteins that are unfolded by damage or mutations (Buneeva and Medvedev, 2018; Mao, 2021). The 19S regulatory particle (RP) sits on one or both sides of the CP and is primarily required for the degradation of structured proteins via mechanically translocating substrates into the degradation chamber of the CP.

The degradation of structured proteins in eukaryotic organisms is predominantly regulated by the ubiquitin-proteasome system (UPS) (Langin and Üstün, 2023; Xu and Xue, 2019). UPS involves prior decoration of substrates with ubiquitin chains via the complex and consecutive actions of E1, E2 and E3 ligase enzymes (Buneeva and Medvedev, 2018; Komander and Rape, 2012; Leestemaker and Ovaa, 2017; Saeki, 2017; Yu and Matouschek, 2017). The ubiquitin chains are recognized by 19S RP substrate receptors regulatory particle non-ATPase (Rpn) 1, Rpn10 and Rpn13, which enable substrate degradation by translocating stretched unstructured regions into the 20S CP channel. Efficient degradation is also dependent on Rpn11 that removes the ubiquitin units during substrate translocation. Beyond binding ubiquitinated substrates, the multimodular Rpn1, Rpn10 and Rpn13 enable transient interactions with ubiquitin-binding shuttle factors, such as RADIATION-SENSITIVE23 (RAD23), and ubiquitin processing enzymes.

A number of bacterial effector proteins target or co-opt the UPS. Among these, SAP05 effectors of bacterial phytoplasma pathogens enable the degradation of structured proteins by the 26S proteasome in a ubiquitin-independent manner (Huang et al., 2021). SAP05 binds both the von Willibrand factor type A (vWA) domain of Rpn10 and the zinc-finger (ZnF) domain of multiple members of two distinct plant transcription factor families, known as SQUAMOSA-PROMOTER BINDING PROTEIN-LIKE (SPL) and GATA BINDING FACTORS (GATAs) (Huang et al., 2021). By hijacking Rpn10, SAP05 mediates the degradation of these TFs, leading to dramatic changes in plant development that includes leaf and stem proliferations, neoteny and increased longevity (Huang et al., 2021), as well as increased plant colonization of phytoplasma insect vectors (Huang and Hogenhout, 2022). Other phytoplasma effectors, known as SAP54/phyllogens, were found to hijack the 26S proteasome shuttle factor RAD23 to mediate the degradation of plant MCM1, AGAMOUS, DEFICIENS, and SRF, serum response factor (MADS) domain transcription factors leading to the induction of leaf-like flowers and other changes in flower development (MacLean et al., 2014), and this also occurs in a ubiquitin-independent manner (Kitazawa et al, 2022). Therefore, phytoplasma effectors appear to directly target ubiquitin receptors or shuttle factors to mediate ubiquitin-independent protein degradation. However, the precise biochemical mechanisms by which these small effector proteins form connections between unrelated proteins to mediate targeted protein degradation in a ubiquitin-independent manner remain uncharacterized.

Here, we determined how SAP05 mediates ubiquitin-independent targeted protein degradation by generating crystal structures of SAP05 in complex with the ZnF domain of SPL5 and with the vWA domain of Rpn10, and validating the ensuing mechanistic model using mutagenesis and protein degradation experiments. We found that SAP05 acts like a molecular glue by bridging ZnF and vWA on opposing surfaces at 1:1 stoichiometries. Furthermore, SAP05 binding to vWA does not appear to cause steric hindrance with other 26S proteasome components and their interactors. Our data show how the bacterial SAP05 effector has evolved as a functional adapter of the 26S proteasome to bypass the canonical UPS cellular proteolysis pathway and enable ubiquitin-independent degradation of structured eukaryotic proteins.

## Results

### Crystal structures of the SAP05 – ZnF_SPL5 and SAP05 – vWA_Rpn10 complexes reveal two distinct binding faces of SAP05

We previously demonstrated that SAP05 of Aster Yellows phytoplasma strain Witches Broom (AYWB) forms a ternary complex with the ZnF domain of SPL5 (ZnF_SPL5) and vWA domain of Rpn10 (vWA_Rpn10) (Huang et al., 2021). To investigate how SAP05 binds these two larger proteins, we determined crystal structures of SAP05 in complex with ZnF_SPL5 and with vWA_Rpn10. We expressed constructs containing SAP05 residues 33 to 135 that correspond to the entire mature part of SAP05 (without the first 32 amino acids encoding the signal peptide that is cleaved off during secretion of the effector), residues 60 to 127 of *Arabidopsis thaliana* SPL5 (accession number AT3G15270) corresponding to the ZnF domain and residues 2 to 193 comprising the vWA domain of *A. thaliana* Rpn10 (accession number AT4G38630) (Supplementary Fig. 1A) in *Escherichia coli*. SAP05 and ZnF_SPL5 were individually expressed and successfully purified at high purity with immobilized metal-affinity chromatography (IMAC) via 6×His tags followed by tag removal and gel filtration (Superdex 75 26/60) (Supplementary Fig. 1B). As vWA_Rpn10 formed aggregates upon purification in the absence of SAP05, vWA_Rpn10 was co-expressed with SAP05 and purified as a complex using the same general method (Supplementary Fig. 1B).

To generate the SAP05 – ZnF_SPL5 complex, we mixed equimolar amounts of purified SAP05 and ZnF_SPL5. We obtained crystals for both complexes, which yielded X-ray data to 2.20 Å resolution for SAP05 – ZnF_SPL5 and to 2.17 Å resolution for SAP05 – vWA_Rpn10. The structure of the SAP05 – ZnF_SPL5 complex was solved via the single-wavelength anomalous diffraction method due to the presence of Zn^2+^ ions bound to ZnF, and that of SAP05 – vWA_Rpn10 was solved with the molecular replacement method using a copy of SAP05 from the SAP05 – ZnF_SPL5 structure and a homology model for vWA_Rpn10 as templates. The details of X-ray data processing and structure solution are described in the materials and methods. The X-ray data collection, refinement, and validation statistics are shown in Table 1.

**Table 1.**
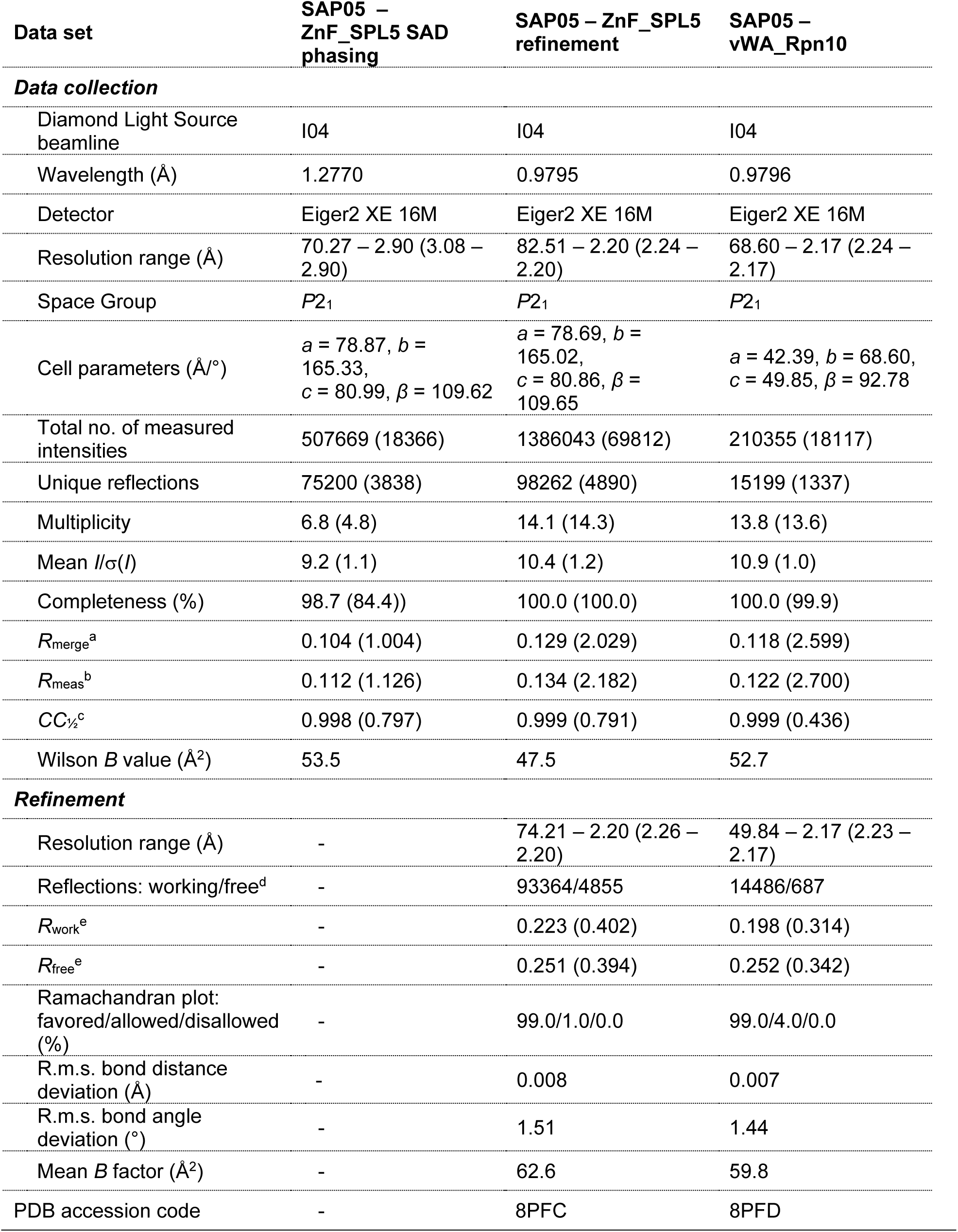
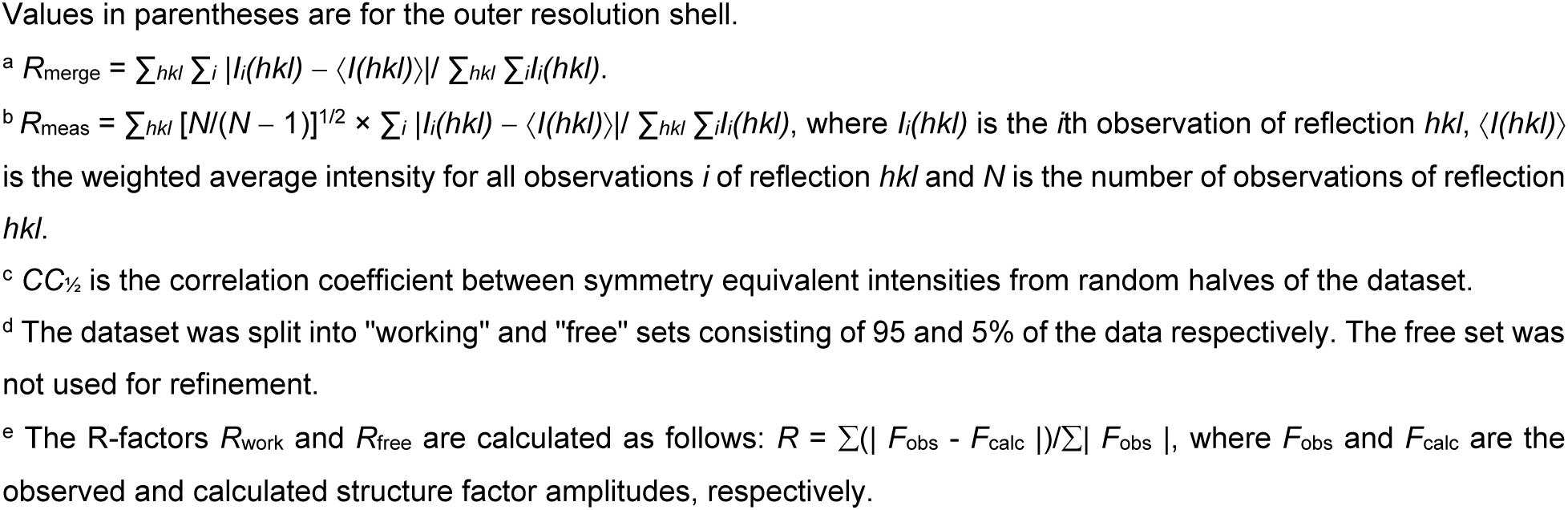
X-ray data collection, processing, and refinement statistics.

| Data set | SAP05 –<br>ZnF_SPL5 SAD<br>phasing | SAP05 – ZnF_SPL5<br>refinement | SAP05 –<br>vWA_Rpn10 |
| --- | --- | --- | --- |
| **Data collection** |  |  |  |
| Diamond Light Source<br>beamline | I04 | I04 | I04 |
| Wavelength (Å) | 1.2770 | 0.9795 | 0.9796 |
| Detector | Eiger2 XE 16M | Eiger2 XE 16M | Eiger2 XE 16M |
| Resolution range (Å) | 70.27 – 2.90 (3.08 –<br>2.90) | 82.51 – 2.20 (2.24 –<br>2.20) | 68.60 – 2.17 (2.24 –<br>2.17) |
| Space Group | *P*2<sub>1</sub> | *P*2<sub>1</sub> | *P*2<sub>1</sub> |
| Cell parameters (Å/°) | *a* = 78.87, *b* =<br>165.33,<br>*c* = 80.99, $\beta$ = 109.62 | *a* = 78.69, *b* =<br>165.02,<br>*c* = 80.86, $\beta$ =<br>109.65 | *a* = 42.39, *b* = 68.60,<br>*c* = 49.85, $\beta$ = 92.78 |
| Total no. of measured<br>intensities | 507669 (18366) | 1386043 (69812) | 210355 (18117) |
| Unique reflections | 75200 (3838) | 98262 (4890) | 15199 (1337) |
| Multiplicity | 6.8 (4.8) | 14.1 (14.3) | 13.8 (13.6) |
| Mean $I/\sigma(I)$ | 9.2 (1.1) | 10.4 (1.2) | 10.9 (1.0) |
| Completeness (%) | 98.7 (84.4)) | 100.0 (100.0) | 100.0 (99.9) |
| $R_{\text{merge}}^a$ | 0.104 (1.004) | 0.129 (2.029) | 0.118 (2.599) |
| $R_{\text{meas}}^b$ | 0.112 (1.126) | 0.134 (2.182) | 0.122 (2.700) |
| $CC_{1/2}^c$ | 0.998 (0.797) | 0.999 (0.791) | 0.999 (0.436) |
| Wilson *B* value (Å<sup>2</sup>) | 53.5 | 47.5 | 52.7 |
| **Refinement** |  |  |  |
| Resolution range (Å) | - | 74.21 – 2.20 (2.26 –<br>2.20) | 49.84 – 2.17 (2.23 –<br>2.17) |
| Reflections: working/free<sup>d</sup> | - | 93364/4855 | 14486/687 |
| $R_{\text{work}}^e$ | - | 0.223 (0.402) | 0.198 (0.314) |
| $R_{\text{free}}^e$ | - | 0.251 (0.394) | 0.252 (0.342) |
| Ramachandran plot:<br>favored/allowed/disallowed<br>(%) | - | 99.0/1.0/0.0 | 99.0/4.0/0.0 |
| R.m.s. bond distance<br>deviation (Å) | - | 0.008 | 0.007 |
| R.m.s. bond angle<br>deviation (°) | - | 1.51 | 1.44 |
| Mean *B* factor (Å<sup>2</sup>) | - | 62.6 | 59.8 |
| PDB accession code | - | 8PFC | 8PFD |

Values in parentheses are for the outer resolution shell.
<sup>a</sup> $R_{\text{merge}} = \sum_{hkl} \sum_i |I_i(hkl) - \langle I(hkl) \rangle| / \sum_{hkl} \sum_i I_i(hkl).$
<sup>b</sup> $R_{\text{meas}} = \sum_{hkl} [N(N-1)]^{1/2} \times \sum_i |I_i(hkl) - \langle I(hkl) \rangle| / \sum_{hkl} \sum_i I_i(hkl),$ where $I_i(hkl)$ is the $i$ th observation of reflection $hkl$ , $\langle I(hkl) \rangle$ is the weighted average intensity for all observations $i$ of reflection $hkl$ and $N$ is the number of observations of reflection $hkl$ .
<sup>c</sup> $CC_{1/2}$ is the correlation coefficient between symmetry equivalent intensities from random halves of the dataset.
<sup>d</sup> The dataset was split into "working" and "free" sets consisting of 95 and 5% of the data respectively. The free set was not used for refinement.
<sup>e</sup> The R-factors $R_{\text{work}}$ and $R_{\text{free}}$ are calculated as follows: $R = \sum(|F_{\text{obs}} - F_{\text{calc}}|) / \sum |F_{\text{obs}}|$ , where $F_{\text{obs}}$ and $F_{\text{calc}}$ are the observed and calculated structure factor amplitudes, respectively.

The structure of SPL5 ZnF resembled the previously determined NMR structures of ZnF domain of *A. thaliana* SPL4 (PDB 1UL4; rmsd = 1.56 Å) and SPL7 (PDB 1UL5; rmsd = 1.76 Å) (Yamasaki et al., 2004). As well, our Rpn10 vWA structure is similar to that of vWA in the spinach 26S proteasome Cryo-EM structure (PDB 8AMZ; rmsd = 0.91 Å) (Kandolf et al., 2022). However, SAP05 did not show any significant structural similarity to experimentally determined structures in the PDB, suggesting a novel architecture.

The SAP05 protein comprises a globular compact structure with five β-strands that form an internal triangular mixed β-sheet core (Fig. 1A, B). β-strand 1 (β1) locates on the long end of the triangle and connects via a loop-helix-loop-helix-loop structure to β2 at the opposite surface near the tip of the triangle. This β-strand is connected to β3 and β4 via loop structures that form the loop-dominated surface of the protein. β4 then connects to a loop-helix-loop structure that runs back to near the long end of the triangle to β5, which runs parallel to β1 (Fig. 1A). Both the SAP05 – ZnF_SPL5 and SAP05 – vWA_Rpn10 structures comprise a 1:1 complex (Fig. 1C, D). The SAP05 residues binding ZnF and vWA are located at opposing surfaces of the effector with the ZnF binding surface comprising the loop-dominated surface (loop surface) and the vWA-binding surface the β-sheet-dominated long end of the triangle (sheet surface) (Fig. 1B, E).

**Fig. 1.**
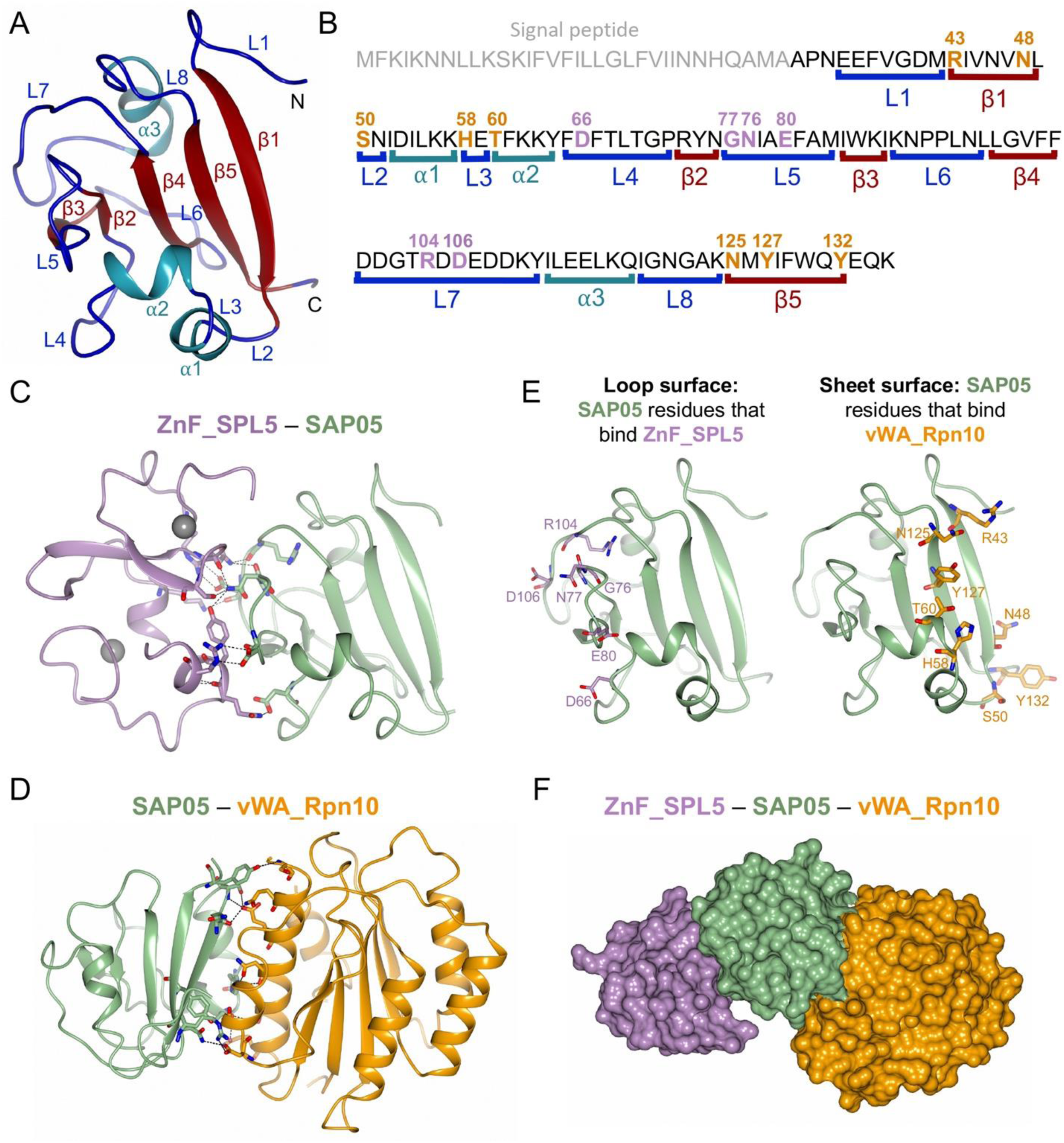
Crystal structures of SAP05 – ZnF_SPL5 and SAP05 – vWA_Rpn10 complexes revealing the bimodular architecture of SAP05. (A) Cartoon model illustrating the fold of the SAP05 effector. α-helices, β-sheets and loops are indicated in cyan, red and blue respectively. (B) Amino acid sequence of SAP05 highlighting the locations of secondary structures (shown in A, colour matched) and ZnF_SPL5 (purple) and vWA_Rpn10 interacting residues (orange) (shown in C-F, colour-matched). ZnF_SPL5, Zinc-finger domain of SPL5 transcription factors. vWA_Rpn10, von Willebrand factor type A domain of Rpn10 ubiquitin receptor. (C) Crystal structure of SAP05 – ZnF_SPL5 complex (PDB 8PFC). The Zn^2+^ ions bound to the two ZnF domains are shown in gray. (D) Crystal structure of SAP05 – vWA_Rpn10 complex (PDB 8PFD). In (C) and (D), the dashed lines indicate the interactions between the residues from both components. (E) Interfaces of SAP05 showing the loop surface (purple) and sheet surface (orange) residues that bind to ZnF_SPL5 and vWA_Rpn10, respectively. (F) Hypothetical ternary structure of ZnF_SPL5 – SAP05 – vWA_Rpn10 obtained by superimposing the crystal structures of ZnF_SPL5 – SAP05 and SAP05 – vWA_Rpn10 complexes.

The SAP05 structures are essentially the same between the SAP05 – ZnF_SPL5 and SAP05 – vWA_Rpn10 complexes (Fig. 1C, D). Moreover, there is no steric hindrance between the ZnF and vWA domains when bound to SAP05, indicating that ZnF and vWA have the capacity to bind SAP05 simultaneously to form a ternary complex (Fig. 1F), consistent with the isolation of ternary complexes containing ZnF, SAP05 and vWA by gel filtration analyses previously (Huang et al., 2021). These data provide evidence that SAP05 has a bimodular architecture with opposite loop and sheet surfaces that enable interactions with ZnF of TFs and vWA of Rpn10, thereby acting as a molecular glue to link SPL and GATA TFs to Rpn10.

### The SAP05 ‘loop surface’ forms electrostatic interactions with the ZnF domain

We further investigated the SAP05 interaction with ZnF_SPL5. The crystal structure contains eight copies of the 1:1 complex in the asymmetric unit (ASU), which are closely similar. In addition to the two structural Zn^2+^ ions within each ZnF_SPL5 domain, a further four Zn^2+^ ions are found in the ASU forming crystal contacts. These involve E80 from four of the eight SAP05 molecules and H82 and E118 from separate neighbouring copies of ZnF_SPL5. The SAP05 loop surface that interacts with ZnF comprises three distinct protruding loops that are separated by β-strands and involves six amino acids of which D66 locates in L4, G76, N77 and E80 in L5 and R104 and D106 in L7 of the SAP05 structure (Fig. 1A, B, E; Fig. 2A). SAP05 binds the two SPL5 ZnF sites, which are held together by Zn^2+^ ions (Fig. 1C; Fig. 2A), and the complex involve 10 connections, with SAP05 amino acids N77 and D106 each binding three amino acids of ZnF_SPL5 and SAP05 D66, G76, E80, and R104 each binding one (Fig. 2A; Supplementary Fig. 2A). The SAP05 – ZnF interaction is dominated by electrostatic interactions of charged and polar residues at the SAP05 loop surface and oppositely charged or polar residues located within ⍺-helices of ZnF_SPL5 (Fig. 2B). To explore the affinity of interaction between SAP05 and ZnF_SPL5, we used isothermal titration calorimetry (ITC). Titration of SAP05 into a solution of ZnF_SPL5 showed an exothermic binding isotherm with a fitted dissociation equilibrium constant (*K*_d_) of 0.45 ± 0.06 µM and stoichiometry of 1:1 (Fig. 2C; Supplementary Fig. 3).

**Fig. 2.**
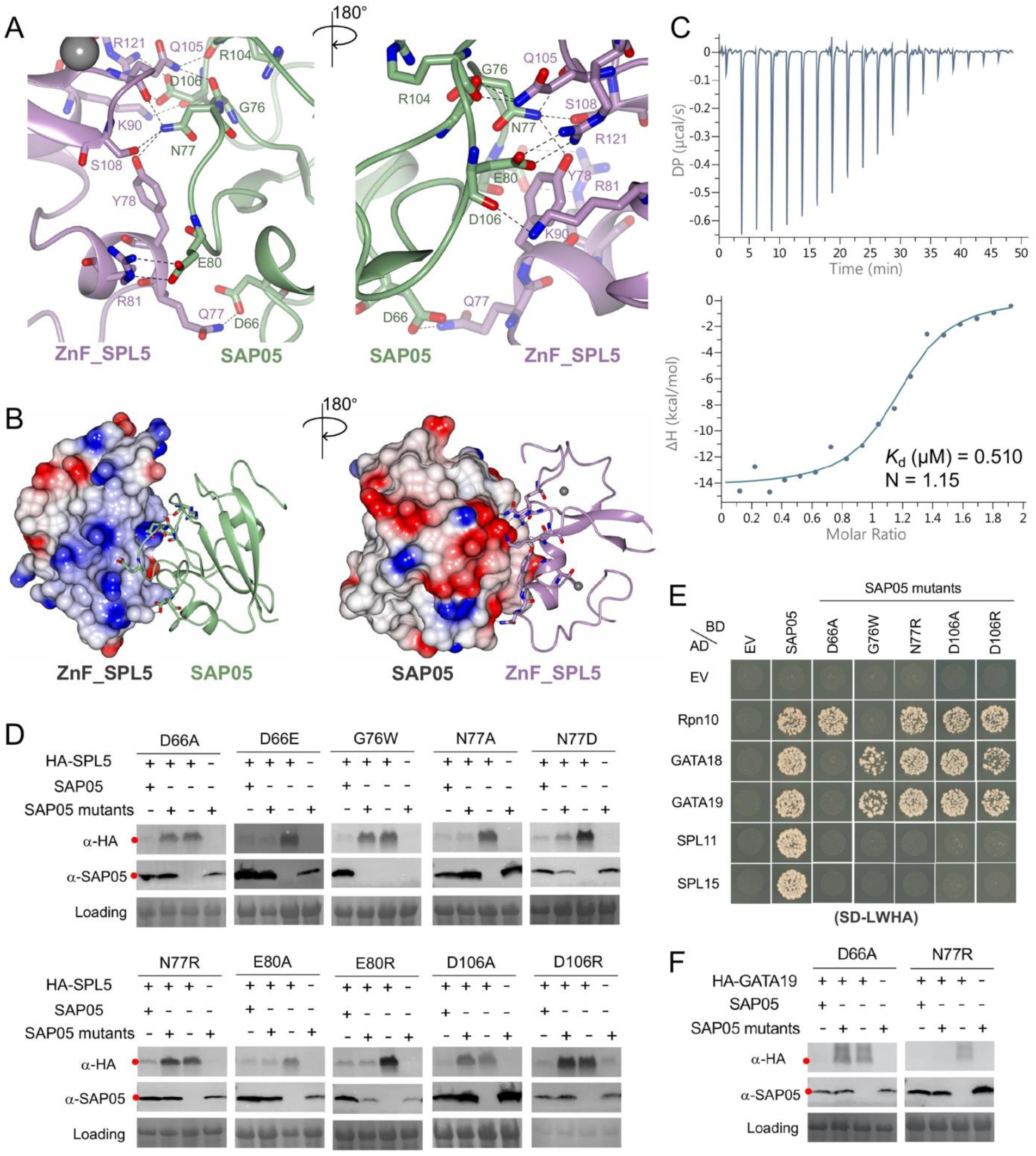
SAP05 loop surface interacts with ZnF_SPL5. (A) Close-up views of SAP05 – ZnF_SPL5 interaction interface from front (left) and back (right). The amino acid residues mediating electrostatic interactions are labelled. The zinc ions are displayed as grey spheres. (B) Electrostatic potential surface view of SAP05 and ZnF_SPL5 during complex formation showing that the interacting interface is predominantly electronegative (red) in SAP05 and electropositive (blue) in ZnF_SPL5. (C) Isothermal titration calorimetry experiment showing direct physical binding of SAP05 and ZnF_SPL5. The top panel shows heat differences upon interaction and the lower panel shows integrated heats of injection (•) and the best fit to a single site binding model using MicroCal PEAQ-ITC analysis software. (D) Western blot analysis of proteasomal degradation of SPL5 in presence of wild-type or mutant SAP05 in *N. benthamiana* leaves. (E) Yeast two-hybrid (Y2H) assay to test interactions of SAP05 and its mutant versions with *A. thaliana* Rpn10 and GATA and SPL TFs. EV, empty vector control. AD, GAL4-activation domain. BD, GAL4-DNA binding domain. SD-LWHA, quadruple dropout medium lacking leucine, tryptophan, histidine and alanine. Yeast growth on -L-W medium is shown in Supplementary Fig. 4A. (F) Western blot analysis for GATA19 degradation in presence of SAP05 D66A or N77R mutants in *N. benthamiana* leaves. (D, F) Red dots indicate the expected sizes of the transiently expressed proteins. HA, Hemagglutinin. Protein loading was visualized using Ponceau S staining.

We generated 10 structure-guided SAP05 mutants by replacing each residue that forms contacts across the interface with a neutral, similarly charged, or oppositely charged residue (Supplementary Fig. 2B). The mutants were investigated for their ability to degrade TFs in *N. benthamiana* leaves using *Agrobacterium*-mediated transient expression assays. SAP05 D66A, G76W, N77R, D106A and D106R failed to degrade SPL5, though G76W was not detected in leaves (Fig. 2D). Yeast two-hybrid (Y2H) assays confirmed that these mutants lost the ability to bind SPL TFs, and except for G76W, retained their affinity for Rpn10 (Fig. 2E; Supplementary Fig. 4). G76W, N77R, D106A and D106R also retained the ability to bind GATA TFs unlike D66A that lost the ability to bind both SPLs and GATAs (Fig. 2E). Consistent with these binding activities in Y2H, D66A did not mediate degradation of GATA19 in leaves whereas N77R did (Fig. 2F). Therefore, D66, N77 and D106 are required for the SAP05 interactions with the ZnF domain of SPLs. Moreover, D66 is also required for SAP05 interactions with GATA TFs. The finding that SAP05 G76W lost binding to both SPLs and Rpn10 suggest that this mutant has more profound structural changes, in agreement with its instability in *N. benthamiana* leaves (Fig. 2D). D66 is part of L4, N77 of L6 and D106 of L7 (Fig. 1A, B; Fig. 2A) indicating that all three loop structures of the SAP05 loop surface are involved in binding ZnF of SPLs (Fig. 1C, E), thereby validating the crystal structure.

Our finding that D66 on one of the loops (L4) is involved in binding GATA prompted us to use AlphaFold-Multimer modeling (Richard et al., 2022) to assess the SAP05 – ZnF_GATA19 complex structure. The AlphaFold model (AFM) of the SAP05 and ZnF_SPL5 complex and their interactions were consistent with the crystal structure (Supplementary Fig. 5A), suggesting a modeling approach could be insightful. We therefore proceeded to predict the SAP05 – ZnF_GATA19 structure using AFM and obtained a high prediction confidence score (Supplementary Fig. 5B). In the model, ZnF_GATA19 interacts with the SAP05 loop surface (Supplementary Fig. 5B). Moreover, SAP05 D66, but not N77, is one of the residues involved in the interaction with GATA19 (Supplementary Fig. 5C), in agreement with the finding that SAP05 N77R degraded GATA19 in *N. benthamiana* leaves, unlike SAP05 D66A (Fig. 2F). SAP05 F65 of loop 4, E80 of loop 5 and D108 of loop 7 were also predicted to mediate interactions with GATA19 (Supplementary Fig. 5C).

Taken together, these data demonstrate that SAP05 interactions with SPL and GATA TFs involves structures of the SAP05 ‘loop surface’. Most SAP05 residues involved in binding SPLs and GATAs do not play a role in SAP05 binding of Rpn10 consistent with the bimodular architecture of SAP05.

### The SAP05 ‘sheet surface’ forms polar interactions with vWA_Rpn10

We then further investigated the role of specific residues involved in the SAP05 – vWA_Rpn10 interface. The SAP05 ‘sheet surface’ comprises two parallel β-strands, β1 and β5, as well as two loops separated by an ⍺-helix (Fig. 1A, B, D, E). The interaction is mediated by eight SAP05 amino acids, including R43 and N48 located on β1, S50 on L2, H58 on L3, T60 on ⍺2, and N125, Y127 and Y132 on β5 of SAP05 (Fig. 3A; Supplementary Fig. 2C). Each of these residues interact with one residue of vWA_Rpn10, except for T60 that interacts with two residues (Fig. 3A; Supplementary Fig. 2C). All vWA_Rpn10 residues that interact with SAP05 locate on ⍺-helices (Fig. 3A). The SAP05 – vWA interaction is largely mediated by polar forces. Thus, the SAP05 ‘sheet surface’ contains rigid secondary β-sheets and ⍺-helix structures that are held in place by hydrogen bonds within SAP05, as opposed to the ZnF-binding loop surface that involves protruding loop structures that are more flexible.

**Fig. 3.**
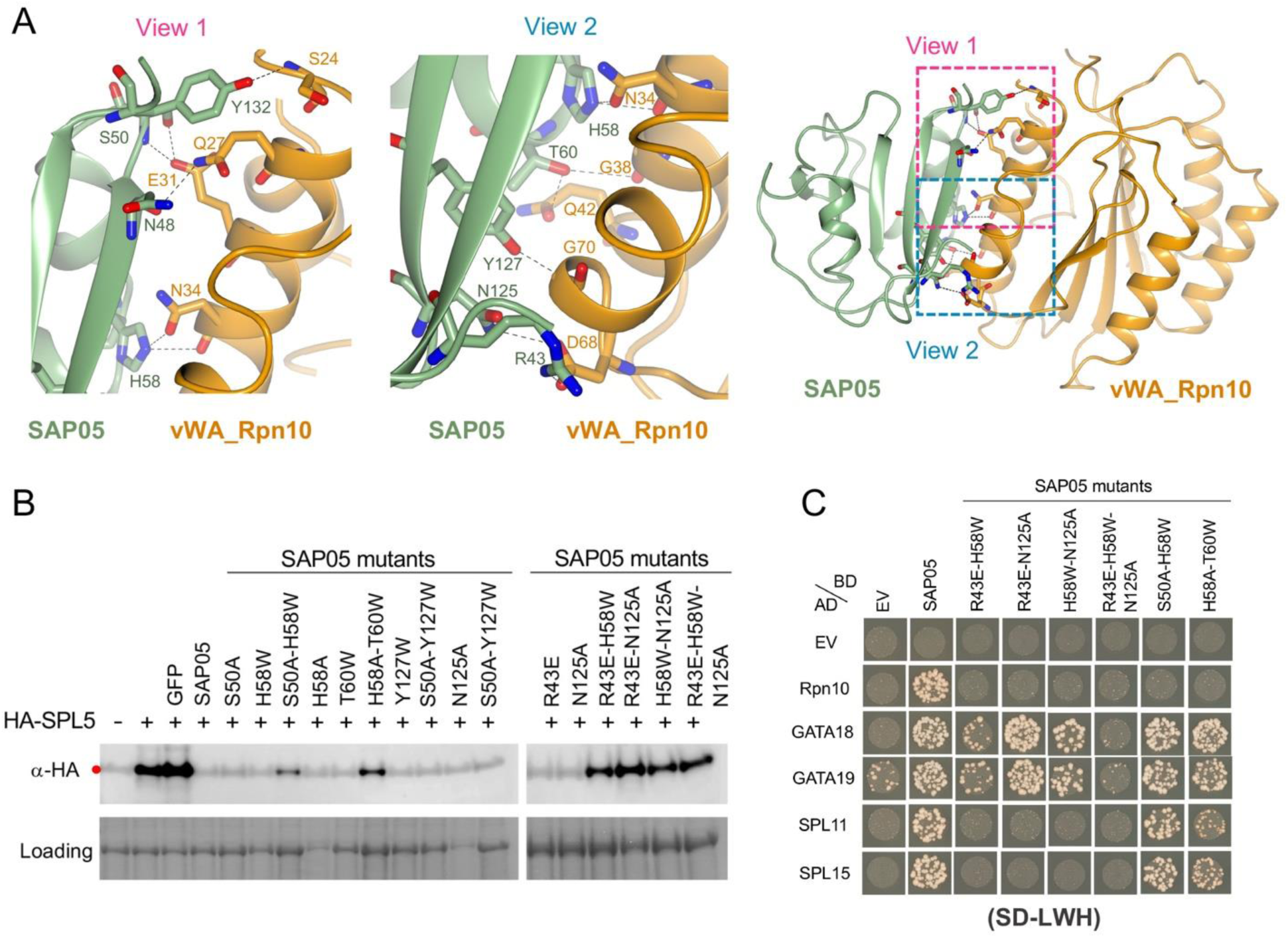
SAP05 β-sheet surface binds to vWA_Rpn10. (A) Stereo views of SAP05 **–** vWA_Rpn10 interaction interface showing the residues involved in complex formation. Right, overview of the interacting interface with two dashed squares displaying the areas for enlarged view. Left, the enlarged view 1 of the top part in the interface. Middle, the enlarged view 2 of the lower part in the interface. (B) Western blot analysis for SPL5 degradation with SAP05 wild-type and mutants in *A. thaliana* protoplasts. GFP, Green fluorescent protein control. HA, hemagglutinin. Protein loading was visualized using Amido Black staining. (C) Y2H assay to test interactions of SAP05 and its mutants with *A. thaliana* Rpn10 and GATA and SPL TFs. EV, empty vector control. AD, GAL4-activation domain. BD, GAL4-DNA binding domain. SD-LWH, triple dropout medium lacking leucine, tryptophan and histidine. Yeast growth on -L-W medium is shown in Supplementary Fig. 4B.

We introduced single amino acid mutations in the SAP05 ‘sheet surface’ by replacing each with neutral, non-polar or oppositely charged residues generating nine single amino acid SAP05 mutants (Supplementary Fig. 2D). All retained the ability to degrade SPL5 in *N. benthamiana* leaves, though some of the SAP05 mutants were not detected in leaves and hence may have high turnover rates (Supplementary Fig. 6). However, several SAP05 double mutants and one triple mutant had reduced or no ability to mediate degradation of SPL5 (Fig. 3B), consistent with their loss of binding to Rpn10 in Y2H (Fig. 3C). SAP05 H58A T60W and S50A H58W retained affinity to SPL and GATA TFs, indicating that these double mutations had minimal impacts on the overall structure of SAP05, unlike the other double mutants that lost affinity to SPLs and the triple mutant that did not bind any of the targets (Fig. 3C). These data indicate that multiple SAP05 residues mediate interactions with vWA, thereby validating the crystal structure. S50, H58 and T60 contribute to this interaction without an obvious impact on SAP05 interactions with the TFs, in line with the bimodular architecture of SAP05.

### Conservation analyses of SAP05 residues involved in SAP05-ZnF, SAP05-vWA interaction reveal dynamic evolutionary patterns

We previously identified SAP05 homologs in phytoplasmas, including examples that bind both SPLs and GATAs, and ones that bind only SPLs or only GATAs (Huang et al., 2021). SAP05 amino acids that interact with SPL5 and vWA are conserved among all or most SAP05 homologs, including D66 that mediates binding with SPL and GATA TFs (Supplementary Fig. 7A). The SAP05 homologs that bind only SPLs (PnWBa, WBDLa and P. mali) and that bind only GATAs (PnWBb, WBDLb) had the most sequence differences in the interacting residues among the SAP05 homologs (Supplementary Fig. 7A). We noticed that regions corresponding to SAP05 residues 67 to 73 (FTLTGPR) that form L4 and connects to β2 in SAP05_AYWB (Fig. 1A, B) were different in sequence among the SPL versus GATA-binding SAP05 homologs (Supplementary Fig. 7A). L4 starts with the conserved F65 and D66 amino acids (Fig. 1A, B; Supplementary Fig. 7A). Given our finding herein that D66 is essential for both SPL and GATA binding and degradation (Fig. 2), we investigated if L4 is involved in determining SAP05-binding specificity for SPLs and GATAs. Swapping corresponding L4 sequences from SAP05 homologs PnWBa and WBDLa to PnWBb and WBDLb resulted in gain of SPL binding, and conversely, swapping these from PnWBb and WBDLb to PnWBa and WBDLa resulted in gain of GATA binding (Supplementary Fig. 7B, C). Therefore, L4 contributes to SAP05 binding specificity to SPLs and GATAs.

Next, we determined in how far interacting residues are conserved among the ZnF domains of SPL TFs. The zinc-binding domain of SPL proteins contains two zinc-binding sites formed by eight conserved Cys or His residues (Yamasaki et al., 2004). SAP05-binding residues locate in both ZnF domains and the majority of these are conserved among the SPL TFs (Supplementary Fig. 8). However, SAP05 D66 interacts with Q77 in SPL5 and this amino acid is not conserved (Supplementary Fig. 8). Given the importance of SAP05 D66 in mediating interactions with both SPL and GATAs, we examined the electrostatic surface in the position of SAP05 D66. In wild-type SAP05, the surface area around D66 is electronegative (Supplementary Fig. 9A, 9C), while the electronegativity of this surface area is reduced in the SAP05 D66A mutant that lost interaction with SPL5 and GATA19 (Supplementary Fig. 9A, 9C). Furthermore, the SAP05 D66E mutant, which is equally charged with negative residue glutamic acid, still degrades SPL5 *in planta* (Fig. 2D). Taken together, these results indicate that the electronegative surface potential contributed by D66 is an important factor for binding SPL and GATA TFs.

Among the SAP05 residues that bind vWA, particularly H58, in combination with S50 or T60, have essential roles (Fig. 3). These locate in L3 that connect ⍺1 and ⍺2 in the SAP05 structure (Fig. 1A, B). SAP05 H58 is conserved among the majority of SAP05 homologs (Supplementary Fig. 7A) and interacts with N34 that is conserved in vWA domains of most Rpn10 homologs (Supplementary Fig. 10). SAP05 S50 interacts with *A. thaliana* vWA E31 that is also an acidic amino (D) in other vWA sequences, and SAP05 T60 with G38 and Q42 that are conserved among plant vWA (Supplementary Fig. 10). The four amino acids locate in the first ⍺-helix (positions 25 - 43) at the N-terminus of *A. thaliana* vWA (Fig. 4; Supplementary Fig. 10).

**Fig. 4.**
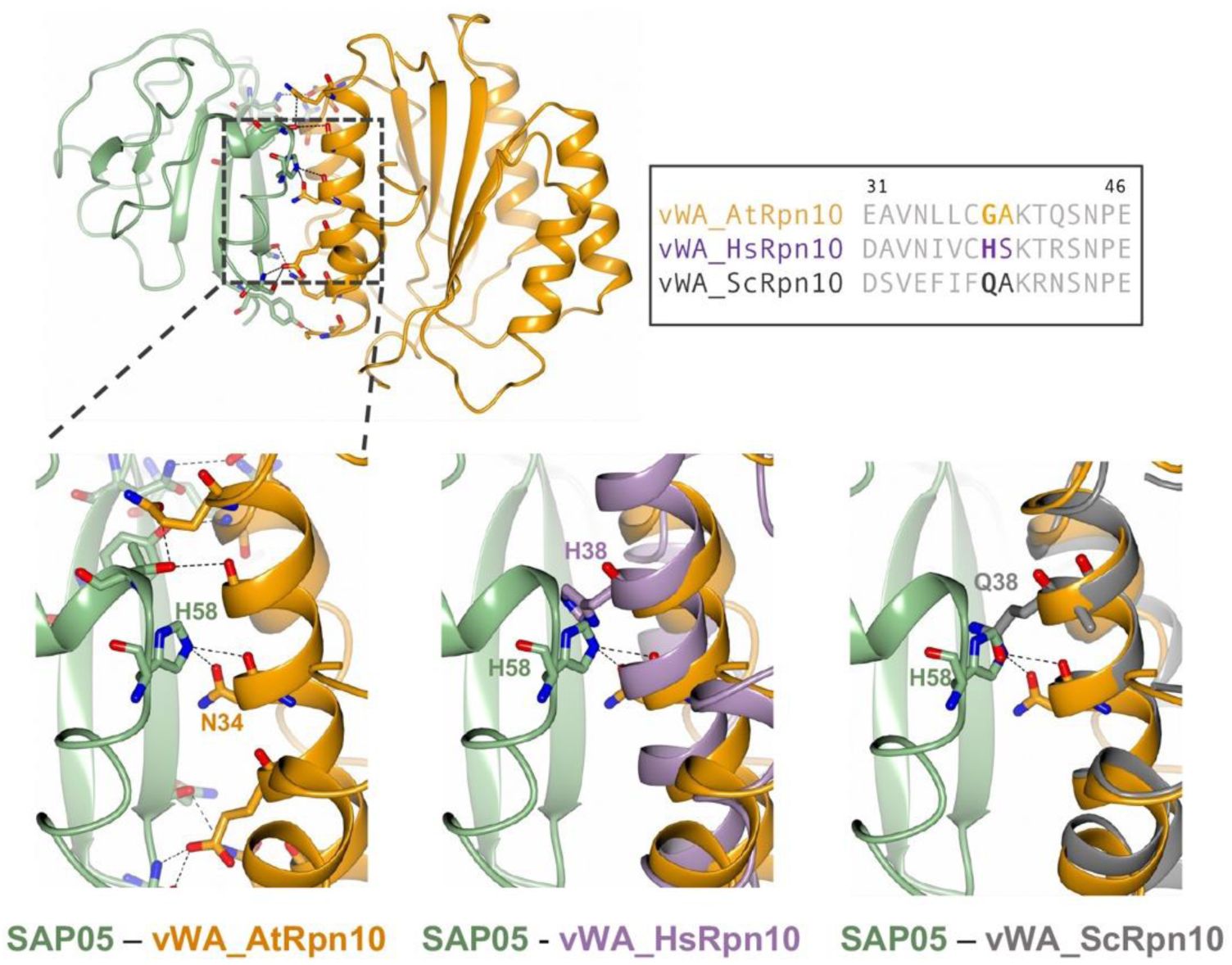
Steric clash at SAP05 loop 3 region prevents SAP05 binding to human and yeast vWA domains. Structure of SAP05 – vWA_AtRpn10 complex is superimposed on the structure of vWA domains from human *Homo sapiens* Rpn10 (PSMD4, PDB 6MSD) or the yeast *Saccharomyces cerevisiae* Rpn10 (PDB 5LN1). Dashed squared box on top left, position of loop 3. Box on top right, sequence alignment of vWA sequences from *A. thaliana* Rpn10 (Uniprot ID: P55034); *H. sapiens* Rpn10 (Uniprot ID: Q5VWC4); ScRpn10, *S. cerevisiae* Rpn10 (Uniprot ID: P38886).

We previously reported that SAP05 does not interact with insect and human Rpn10 (the latter is also known as PSMD4), and that replacing AtvWA G38 and A39 with human H38 and S39 respectively prevented SAP05 binding (Huang et al., 2021). Multiple sequence alignment of plant and animal vWA domains showed high conservation with a few amino acid differences (Supplementary Fig. 10). Structural superimposition of SAP05 interactions with vWA domains of plant and PSMD4 revealed that SAP05 L3 clashes with H38 of PSMD4 precluding a SAP05-PSMD4 interaction (Fig. 4). Similarly, Q38 of vWA of yeast Rpn10 clashes with L3 of SAP05 precluding SAP05 from binding yeast Rpn10 (Fig. 4). This corroborates data presented herein that H58 of the SAP05 L3 region plays an essential role in the SAP05 interaction with *A. thaliana* vWA.

### Positioning of SAP05 complex on the 26S proteasome points to a TPD mechanism

Rpn10 is positioned in the 19S RP where it forms an important component in the interface of the lid and base structure (Glickman et al., 1998). The cryo-EM structure of the spinach (*Spinacia oleracea*) 26S proteasome was recently resolved (Kandolf et al., 2022). The vWA residues that interact with 26S protesome are conserved in the spinach *Spinacia oleracea* and *A. thaliana* Rpn10 homologs (Supplementary Fig. 11). The structural superposition of SAP05 – vWA_Rpn10 complex onto the spinach 26S proteasome did not reveal obvious steric clashes with proteasome components (Fig. 5A). Notably, SAP05 interacts with two parallel ⍺-helices that locate on an area of vWA that does not interact with the 19S subunit (Fig. 5B). This suggests that SAP05 has minimal disadvantageous effects on Rpn10 interactions with the 19S RP. However, the ZnF_SPL5 of the ternary structure sterically clashes with two ⍺-helices that protrude from the 26S proteasome (Supplementary Fig. 12). We found that these helices are derived from the flexible N-terminal coiled-coil (CC) domains of the spinach homologs of *A. thaliana* Rpt4 and Rpt5 (Supplementary Fig. 12), which are part of the 6 AAA+ ATPase (Rpt1-6) subunit ring that sits on top of the 20S CP. The CC domains of these Rpt proteins form an important role in physically connecting substrate recruitment and processing (Snoberger et al., 2018), and may therefore be involved in ZnF_SPL5 and degradation. We propose that interaction of SPL5 with the CC dimer may induce the degradation of the transcription factor.

**Fig. 5.**
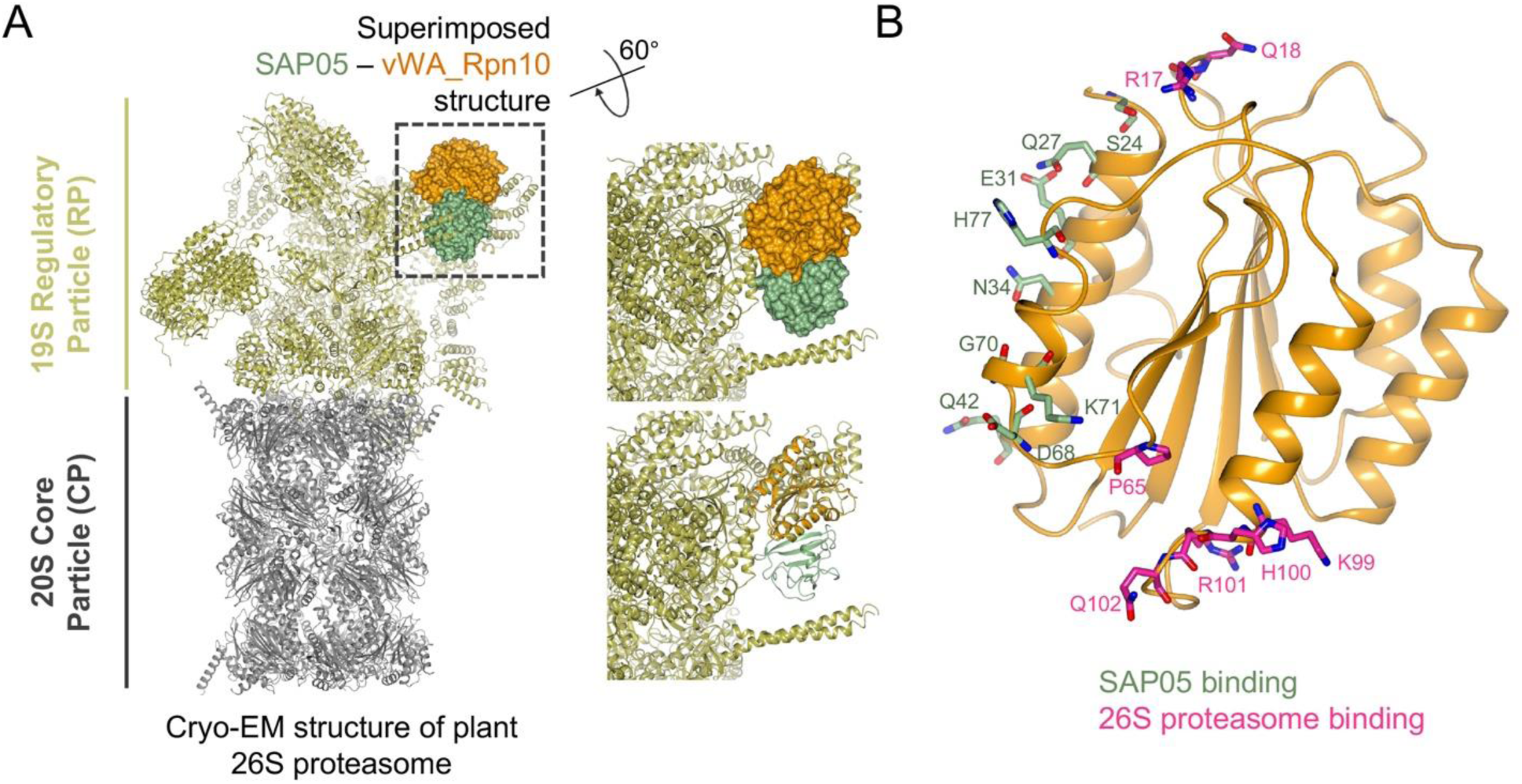
SAP05 interaction with vWA_Rpn10 domain has no hindrance on plant 26S proteasome. (A) Structural superimposition of SAP05 – vWA_Rpn10 structure on the spinach 26S proteasome (PDB 7QVE and PDB 8AMZ). Dashed square box shows the parts with superimposition and is enlarged to show the surface (right top) and cartoon (right bottom) view of SAP05 – vWA_Rpn10 complex. (B) Position of vWA_Rpn10 residues interacting with SAP05 (green) and subunits of the 26S proteasome (pink).

## Discussion

The crystal structure data reported herein demonstrates that the 12.3 kDa bacterial effector SAP05 has a globular structure with an internal β-sheet core and interconnecting loops and ⍺-helices. One interaction surface is dominated by rigid β-strands (sheet surface) and the opposite surface by flexible loop structures (loop surface). The rigid sheet surface binds the vWA domain of plant Rpn10 and cannot bind vWA domains of Rpn10 homologs of organisms other than plants, even though the vWA domains are highly conserved among Rpn10 homologs. The flexible loop surface is however capable of binding multiple transcription factors of the distinct SPL and GATA TF families via their zinc-finger domains. Therefore, SAP05 appears to have an optimal configuration to act as a molecular bridge capable of connecting a conserved proteasome component to multiple members of two TF families.

Rpn10 vWA has an important function as its deletion causes lethality or severe growth deficiencies in plants, vertebrates and human cell lines (Fatimababy et al., 2009; Hamazaki et al., 2023; Rani et al., 2012). In *rpn10* null mutants, deficiencies may be restored upon addition of the vWA domain of Rpn10 and RAD23, and Rpn10 and RAD23 act redundantly (Elsasser et al., 2004; Verma et al., 2004). Like Rpn10, RAD23 is a reversible component of the 26S proteasome and bind ubiquitinated substrates (Elsasser et al., 2004). Intriguingly, the phytoplasma SAP54/phyllogen family of effectors hijack RAD23 and mediates plant proteins, the MADS-box TFs for degradation, leading to plant developmental changes (Kitazawa et al., 2022; MacLean et al., 2014). SAP54 bind the RAD23 ubiquitin-associated (UBA) that are shown to covalently bind ubiquitin moieties of ubiquitinated substrates (Chen et al., 2001; Saeki, 2017). Therefore, at least two distinct phytoplasma effector families mediate degradation of plant TFs in ubiquitin-independent manner using reversible components of the 19S RP.

The vWA domain locates at the N-terminus of Rpn10, and the C-terminal part of Rpn10 consists of single-helix ubiquitin interaction motifs (UIMs) connected by a flexible region that bind ubiquitin chains (Sakata et al., 2012; Wang et al., 2005). Due to the flexible C-terminus, only the vWA domain is visible in cryo-EM crystal structures of 26S proteasomes and locates in a central location of the 19S RP, at the interface of its base and lid (Chen et al., 2020). There is no evidence that SAP05 interferes with 26S proteasome activity (Huang et al. 2021) and structural information generated herein shows that SAP05 binds to a region of vWA that does not interact with components of the 19S RP. Therefore, SAP05 appears to have evolved to optimally bind plant Rpn10 and is positioned in a central location of the 19S RP.

Our findings indicate that the placement of the SAP05 complex on the 26S proteasome suggests a TPD mechanism. While the SAP05 – vWA_RPN10 complex does not appear to cause significant steric hindrance, the ZnF_SPL5 within the ternary structure appears to clash sterically with the ⍺-helices of Rpt4 and Rpt5, as evidenced in the structural model of the spinach 26S proteasome (Kandolf et al., 2022). The coiled-coil (CC) domains of these two Rpt proteins, along with the domains of four others, dimerize to create three CCs (Rpt1/2, Rpt6/3, and Rpt4/5 CCs). These CCs play a vital role in 26S proteasome activity by physically linking substrate recruitment and processing to the unfolding machinery, ultimately leading to substrate degradation (Matyskiela et al., 2013). Furthermore, conformational changes within the CCs are essential for transitioning the Rpt1-6 ATPase ring between resting and active stages (Snoberger et al., 2018), the latter of which involves a widening of the central pore to allow substrate entry into the core of the 20S CP (Matyskiela et al., 2013). The Rpt4/5 CC also binds to Rpn10 upon substrate binding (Matyskiela et al., 2013). As such, when the ternary complex is accommodated on the 26S proteasome, the interaction of SPL5 with the CC dimer could potentially instigate the switch of the proteasome to the active stage, leading to the degradation of the transcription factor.

We investigated the importance of residues mediating the SAP05 – ZnF_SPL5 and SAP05 – vWA_Rpn10 interfaces. We found that mutations of single amino acids in SAP05 disrupted the SAP05 – ZnF_SPL5 interaction. SAP05 mutant D66A is impaired in the degradation of both SPL5 and GATA19, showing a potential to be further engineered for a useful tool to degrade any protein without causing plant development problems.

In contrast, multiple amino acid mutations are needed to impair SAP05 interaction with vWA_Rpn10. Even in the double SAP05 H58A T60W and S50A H58W double mutants the ability to mediate degradation has not fully disappeared. This suggest that the SAP05 – vWA interaction is robust. It is striking that SAP05 itself is not degraded, and is highly stable in plant cells, despite its association with the 26S proteasome and its function in degrading substrates that it directly interacts with. However, the study herein revealed that SAP05 derivatives with mutations in the SAP05 – vWA_Rpn10 interaction surface are often unstable in plant cells. An explanation of these results is that interactions with vWA of some of the SAP05 single amino acid mutants is weaker than wild type and this leads to SAP05 being dragged down along with ZnF when the latter is pulled into the core of the 20S CP. We were unable to test this with ITC experiments, because purified vWA on its own (without SAP05) forms aggregates. Nonetheless, results so far suggest that SAP05 stability is linked to its association with Rpn10.

In summary, given that SAP05 displayed no notable structural similarity to any experimentally confirmed structures in the Protein Data Bank (PDB), we suggest that this effector protein possesses a previously unknown molecular architecture. This bimodular architecture of SAP05 enables it to connect host proteins to the Ubiquitin Proteasome System (UPS), thereby circumventing the canonical targeted protein degradation pathway. Impaired functioning of the UPS is linked with a multitude of diseases. However, given the pivotal role of the UPS in orchestrating an array of cellular processes, the SAP05 molecular system presents opportunities for the development of innovative therapeutics. Notably, Rpn10 has been recognized as a potential therapeutic target (Du et al., 2023). Furthermore, the 26S proteasome has been leveraged to create novel therapeutics such as PROTACs (PROteolysis TArgeting Chimeras), which are small molecules that recruit E3 ligases for ubiquitination of substrates marked for degradation (Lu et al., 2021). Several PROTACs and similar systems have shown promising results and are currently advancing through clinical trials. However, their reliance on recruiting E3 ligases has led to challenges associated with side effects and resistance. Since SAP05 does not hamper the ability of the 26S proteasome to degrade substrates, this effector emerges as a prime candidate for engineering a new type of degraders that operate independently of E3 ligases. The structural work presented in this study provides a springboard for bioengineering a new class of degraders that do not depend on E3 ligases.

## Materials and Methods

### Protein production and purification

#### Gene cloning, expression, and protein purification for crystallization and *in vitro* studies

For crystallization of SAP05 – ZnF_SPL5 complex, DNA encoding mature SAP05 excluding signal peptide (Ala33-Lys135) and DNA encoding the ZnF domain of SPL5 (Ser60-Leu127) were separately subcloned to pOPINF vector for N-terminal 6×His tags (Berrow et al., 2007) using In-fusion cloning method (Benoit et al., 2006; Bird et al., 2014). The constructs were transformed individually into *E. coli* BL21 (DE3) competent cells. Bacterial cultures were grown in LB media (containing 50 µg/mL ampicillin) at 37 °C to an OD_600_ around 0.5 followed by induction with 1 mM Isopropyl-b-D-thiogalactoside (IPTG) at 16 °C overnight with shaking at a speed of 220 rpm. Cell pellets were resuspended in buffer A1 (50 mM Tris-HCl, 50 mM glycine, 0.5 M NaCl, 20 mM imidazole, 5% glycerol, pH 8.0) followed by sonication lysis at 40% amplitude, 5 sec on / 10 sec off pulse for 30 min on ice. Cell debris was removed by centrifuge at 20000 g for 20 min. Purification of the proteins was performed using an ÄKTA Xpress purification system comprising initial capture with metal affinity chromatography (IMAC) using Ni-NTA column, which elutes proteins with buffer B1 (50 mM Tris-HCl, 50 mM glycine, 0.5 M NaCl, 0.5 M imidazole, 5% glycerol, pH 8.0), followed by gel filtration with the column Superdex 75 26/600 in buffer A4 (20 mM HEPES, 0.15 M NaCl, pH 7.5). Fractions with the elution peaks were assessed by SDS-PAGE gels, pooled and treated with HRV-3C protease overnight at 4°C. Afterwards, digested protein samples were passed through Ni-NTA column to remove cleaved 6×His tags. Untagged samples were assessed with SDS-PAGE gels, pooled and concentrated with 3 kDa cut-off vivaspin concentrators, followed by a second round of gel filtration with Superdex 75 26/600 in buffer A4. Eluted samples were assessed with SDS-PAGE gels. Afterwards, purified proteins were mixed together in equimolar ratio, followed by gel filtration chromatography. Eluted fractions were assessed by SDS-PAGE, pooled and concentrated to a final concentration of 15 mg/mL, for crystallization studies.

For crystallization of SAP05 – vWA_RPN10 complex, the mature coding sequence of SAP05 (Ala33-Lys135) was cloned to pOPINF vector, vWA_RPN10 (Val2-GLy193) was initially cloned to pOPINM to make MBP-vWA fusion cassette. MBP-vWA was then amplified and ligated to pOPINA to remove the 6×His tag. The final constructs were co-transformed into *E. coli* BL21(DE3) competent cells. Proteins were co-expressed and co-purified using the IMAC and gel filtration as mentioned above. Eluted fractions from the complex were assessed from gel filtration peaks and SDS-PAGE. Afterwards, the eluted complex was subjected to HRV-3C protease cleavage overnight at 4°C followed by Ni-NTA column to remove tags. After a second round of gel filtration followed by assessment with SDS-PAGE, the purified complexes were pooled and concentrated to 15 mg/mL for crystallization studies.

For *in vitro* experiments, purified proteins were concentrated to ∼10 mg/mL via 3 kDa cut-off vivaspin concentrators for subsequent analysis. Detection of proteins on SDS-PAGE gels was performed using Instant Blue staining solution (Abcam). Data of gel filtration traces were processed and plotted using ggplot2 in R (Wichham, 2011).

#### Protein crystallization, structure determination, and refinement

Crystallization screens were set up in sitting-drop vapor diffusion format in MRC2 96-well crystallization plates with drops comprised of 0.3 μL precipitant solution and 0.3 μL of protein and incubated at 293 K. All crystals were cryoprotected in the crystallization solution supplemented with 25% (v/v) glycerol and mounted in Litholoops (Molecular Dimensions) before flash-cooling by plunging into liquid nitrogen. X-ray data were recorded on beamline I04 at the Diamond Light Source (Oxfordshire, UK) using an Eiger2 XE 16M hybrid photon counting detector (Dectris), with crystals maintained at 100 K by a Cryojet cryocooler (Oxford Instruments). Diffraction data were integrated and scaled using DIALS (Winter et al., 2018) via the XIA2 expert system (Winter, 2010) then merged using AIMLESS (Evans and Murshudov, 2013). The majority of the downstream analysis was performed through the CCP4i2 graphical user interface (Winn et al., 2011). Data collection statistics are summarized in Table 1.

The SAP05 – ZnF_SPL5 complex crystallized from 0.1 M MES pH 6.5, 25% (w/v) PEG 6000 in space group *P*2_1_, with approximate cell parameters of a = 78.7, b = 165.0, c = 80.9 Å, *β* = 109.7°. All X-ray data were collected from a single crystal, initially recorded at a wavelength of 0.9795 Å (2 × 360° passes) and processed to 2.2 Å resolution, and then at a wavelength of 1.2770 Å (1 × 360° pass), this being close to the *K* X-ray absorption edge for zinc. The latter data set was processed to 2.4 Å resolution and enabled structure solution via the single-wavelength anomalous diffraction method using the CRANK2 pipeline (Skubak et al., 2018), due to the presence of Zn^2+^ ions bound to ZnF_SPL5. This produced a partial model corresponding to eight copies of a 1:1 complex of SAP05-ZnF_SPL5 in the crystallographic asymmetric unit, giving an estimated solvent content of 59%, with each copy of ZnF_SPL5 containing two Zn^2+^ ions. The model was completed through several iterations of model building in COOT (Emsley et al., 2010) and restrained refinement in REFMAC5 (Murshudov et al., 2011) against the higher resolution data set at 2.2 Å resolution. Refinement and validation statistics are summarized in Table 1.

The SAP05 – vWA_Rpn10 complex crystallized from 0.1 M Sodium HEPES pH 7.5, 10.7 % (w/v) PEG 4000 in space group *P*2_1_, with approximate cell parameters of a = 42.4, b = 68.6, c = 49.9 Å, *β* = 92.8°. X-ray data were recorded from a single crystal at a wavelength of 0.9796 Å (2 × 360° passes) and processed to 2.17 Å resolution. Analysis of the likely composition of the asymmetric unit suggested that it contained one copy of a 1:1 complex of SAP05-vWA_Rpn10, giving an estimated solvent content of 44%. The structure was solved via molecular replacement using PHASER (McCoy et al., 2007). One copy of SAP05 was taken from the above SAP05-ZnF_SPL5 complex as the first template, and the second template was derived from a homology model of vWA_Rpn10 produced by Swiss-Model (Waterhouse et al., 2018) and based on PDB entry 5VFT (Zhu et al., 2018). The model was completed through several iterations of model building in COOT and restrained refinement in REFMAC5. Refinement and validation statistics are summarized in Table 1. All structural figures were prepared using CCP4mg (McNicholas et al., 2011).

### *In vitro* Protein-Protein Interaction study

#### Isothermal titration calorimetry (ITC)

ITC experiments were performed with MicroCal PEAQ-ITC instrument (Malvern, UK). The data were recorded at 25°C using 20 mM HEPES, pH 7.5, 150 mM NaCl buffer. The calorimetric cell was filled with 20 μM ZnF_SPL5 and titrated with 200 μM SAP05 from the syringe. A single injection of 0.4 μL was followed by 19 injections of 2 μL each. Injections were made at 150 sec interval with a stirring speed of 750 rpm. Each experiment was repeated three times. The raw data were integrated and fitted to a one-site binding model using the built-in software of MicroCal PEAQ ITC (Bastos and Velazquez-Campoy, 2021; Zambelli, 2019).

### SAP05 mutation generation, cloning, yeast two-hybrid (Y2H) assay and degradation assay

#### SAP05 mutation generation and cloning

SAP05 mutants were generated by overlap PCR (Nelson and Fitch, 2011) or directly synthesized using gBlock from Integrated DNA Technologies company (IDT). Mutations used for different assays were codon optimized and cloned to different vectors.

#### Yeast two-hybrid assay (Y2H)

The coding sequences of SAP05 or SAP05 mutants excluding signal peptides were amplified and ligated into Gateway vector pDEST-GBKT7 (BD). Full length sequences of *A. thaliana* Rpn10, two SPLs (SPL11 and SPL15), and three GATAs (GATA18, GATA19 and GATA25) were amplified and cloned into vector pDEST-GADKT7 (AD) using Gateway cloning methods (Hartley et al., 2000; Liang et al., 2013). Constructs used to test protein-protein interactions were co-transformed into the yeast *Saccharomyces cerevisiae* strain AH109 using the Matchmaker Gold yeast two-hybrid system (Clontech). Empty pDEST-GBKT7 and pDEST-GADKT7 vectors were used as negative control. Yeast growth was assessed on solid double dropout medium lacking leucine and tryptophan (SD-LW) which indicates the presence of both AD and BD constructs. Interactions between AD and BD fusion proteins were screened on selective dropout medium; triple dropout medium lacking leucine, tryptophan and histidine (SD-LWH) or with the addition of 3-amino-1,2,4-triazole (3-AT) at a final concentration of 10 mM when self-activation was observed or to improve selection stringency, and quadruple dropout medium lacking leucine, tryptophan, histidine and alanine (SD-LWHA). Yeast plates were kept in 28°C growth chambers for 5 days before imaging.

#### Degradation assay in *N. benthamiana*

The coding sequences of SAP05 or SAP05 mutants (excluding signal peptides) were amplified and ligated into Gateway vector pB7WG2. Full length coding sequence of AtSPL5 were amplified and tagged with 3×HA at N-terminal, then ligated into Gateway vector pB7WG2. After sequencing, the constructs were separately transformed to *Agrobacterium tumefaciens* strain GV3101 and plated on LB solid medium containing rifampicin, gentamicin and spectinomycin and grown at 28 °C for 24-48 h. Then colonies were picked and checked by PCR using plasmids extracted from overnight liquid culture and gene-specific primers. Positive colonies were grown at 28 °C overnight, harvested, and resuspended in infiltration buffer (10 mM MgCl_2_, 10 mM MES, pH 5.6), supplemented with 100 μM acetosyringone. Appropriate combinations of above constructs together with pCB301-P19 were mixed at an OD_600_ of 0.5 per construct and infiltrated to the abaxial surface of 4-week-old *N. benthamiana* leaves using 1 ml needleless syringe. Infiltrations were conducted on randomly selected leaves, with triplicates consisting of three leaves each. To minimize variation, each single leaf was infiltrated with four combinations: SAP05 only, one SAP05 mutant only, SAP05 plus HA-SPL5, and SAP05 mutant plus HA-SPL5. After 3 days post-infiltration, the infiltrated leaves were detached, and the infiltrated areas were harvested for total protein extraction.

To detect SPL5 and SAP05, western blots were performed as follows. Total protein extracts were separated on 4-20% gradient gels (Bio-Rad) and transferred to 0.22 μm PVDF membranes using the Bio-Rad mini-PROTEAN Electrophoresis system. Protein loading was visualized with Ponceau S staining solution (Thermo Scientific) and washed off with water. Membranes were then blocked by 5% (w/v) milk powder in Tris-buffered saline (TBS) and 0.1% (v/v) Tween-20 for 2 h at room temperature. Membranes were then overnight incubated at 4°C with the anti-HA primary antibody (OptimAb HA. 11, Eurogentec), which was raised from mouse serum, at a ratio of 1:2000 dilution. Afterwards, the membrane was probed with Alkaline-Phosphatase-conjugated anti-mouse secondary antibody (Thermo Scientific) for 1 h at room temperature and imaged after incubation with NBT/BCIP substrate solution (Thermo Scientific). Membranes were washed after detection, blocked in the same conditions as previously described and incubated overnight at 4°C with rabbit raised anti-SAP05 antibody at a 1:5000 ratio (Huang et al., 2021). The membrane was then probed with HRP conjugated anti-rabbit secondary antibody (Sigma) and imaged with Immobilon Western Chemiluminescent HRP Substrate (Sigma).

#### Degradation assay in *A. thaliana* protoplast

*A. thaliana* (Col-0) mesophyll protoplast isolation and transformation were carried out as previously reported (Yoo et al., 2007). Briefly, protoplasts from the mesophyll cells were isolated from leaves of 3-4 week-old plants which were grown under short-day (10 h light/14 h dark) conditions at 22°C. For transfection, 300 μL of fresh protoplast solution (400,000/mL) was co-transformed with 12 μg high quality plasmids for each construct using PEG-calcium method. Transfected protoplasts were incubated at 22°C for 16 h in dark. Following which, total protein was extracted and examined via western blots using HRP-conjugated anti-mouse secondary antibody (Sigma). Protein loading was visualized with Amido Black Staining Solution (Sigma).

### Homology and structural analysis

#### Homology analysis

Sequences of SAP05 homologs from different phytoplasma strains, *A. thaliana* SPLs and Rpn10 from different organisms were aligned using MUSCLE algorithm on Phylogeny.fr web server (http://www.phylogeny.fr/index.cgi; Dereeper et al., 2008). Graphical representation and editing were performed with MEGA11 software.

#### Structural analysis

Structural predictions were conducted using AlphaFold v2.3.2 (Jumper et al., 2021) for single protein and AlphaFold-Multimer v3 (Richard et al., 2022) for protein complexes. Analysis of predicted structure was performed with CCP4mg (McNicholas et al., 2011).

#### Data availability

Crystal structure date have been deposited to the Protein Data Bank (PDB ID code 8PFC and 8PFD). All other data are presented in the article and supporting information.

## Acknowledgements

This work was supported by the Human Frontier Science Program (grant number: RGP0024/2015), the UK Research and Innovation (UKRI) Engineering and Physical Sciences Research Council (grant number: EP/X024415/1), start-up funding from the CAS Center for Excellence in Molecular Plant Sciences, the Natural Science Foundation of Shanghai (grant number 23ZR1470300), and the UKRI BBSRC (Grant BBS/E/J/000PR9797) with additional support from the John Innes Foundation and the Gatsby Charitable Foundation. We acknowledge Diamond Light Source for access to beamline I04 under proposal MX25108. We thank Julia Mundy, Biophysical Analysis and Protein Crystallography platform at the John Innes Centre (JIC) for their support with ITC, protein crystallization, and X-ray data collection.

## Supplementary Figures

**Supplementary Fig. 1.**
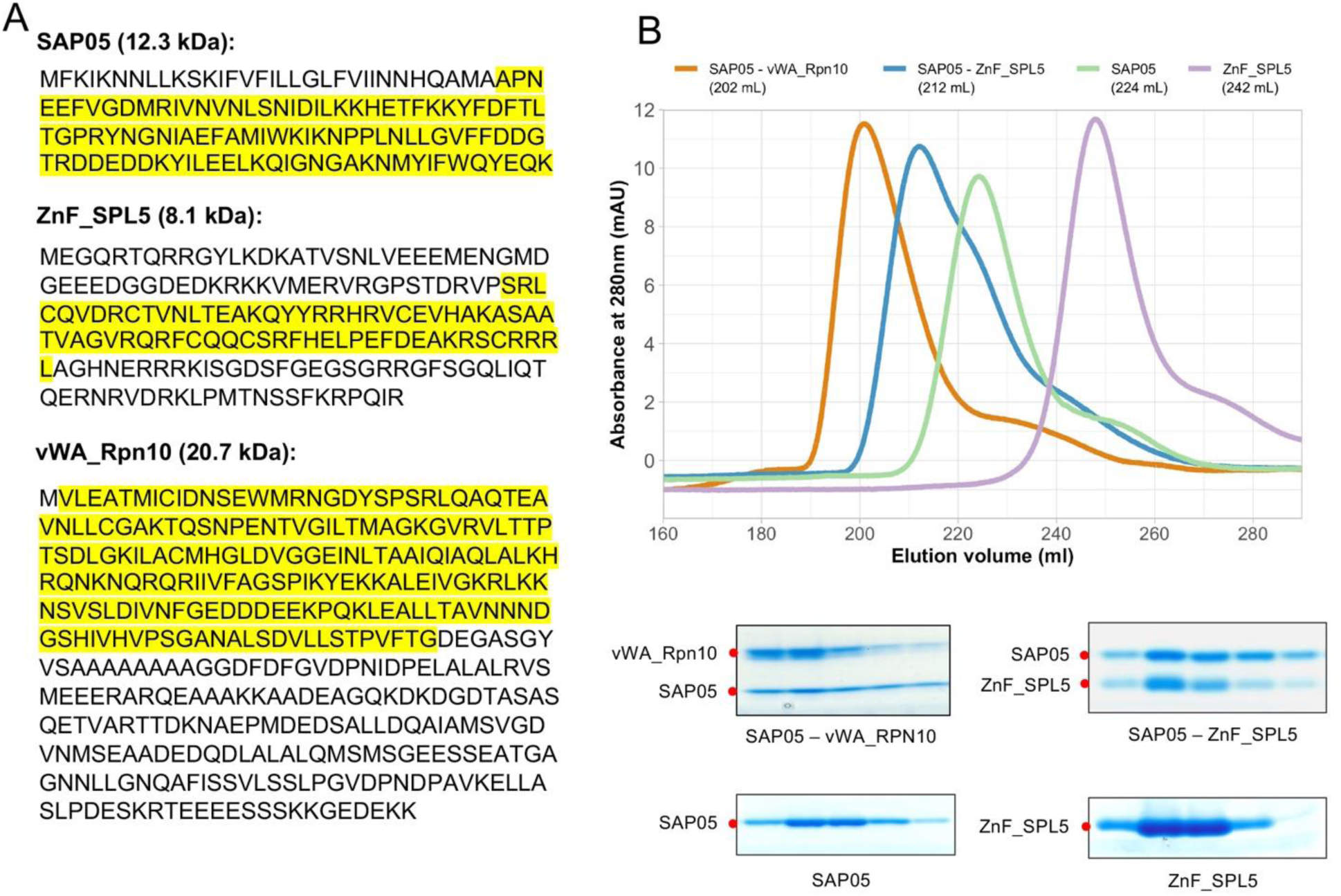
Expression and purification of SAP05 – ZnF_SPL5 and SAP05 – vWA_RPN10 complexes for crystallization. (A) The amino acid sequences of mature SAP05 (without signal peptide) and of ZnF_SPL5 and vWA_Rpn10 utilized for protein expression are highlighted in yellow on full-length proteins and their molecular weights are indicated in brackets. (B) Gel filtration chromatogram (top) and SDS-PAGE (bottom) analyses of peak fractions of purified SAP05, ZnF_SPL5, SAP05 – ZnF_SPL5 and SAP05 – vWA_Rpn10 complexes. Red dots indicate the expected size of protein and protein complexes in the gels.

**Supplementary Fig. 2.**
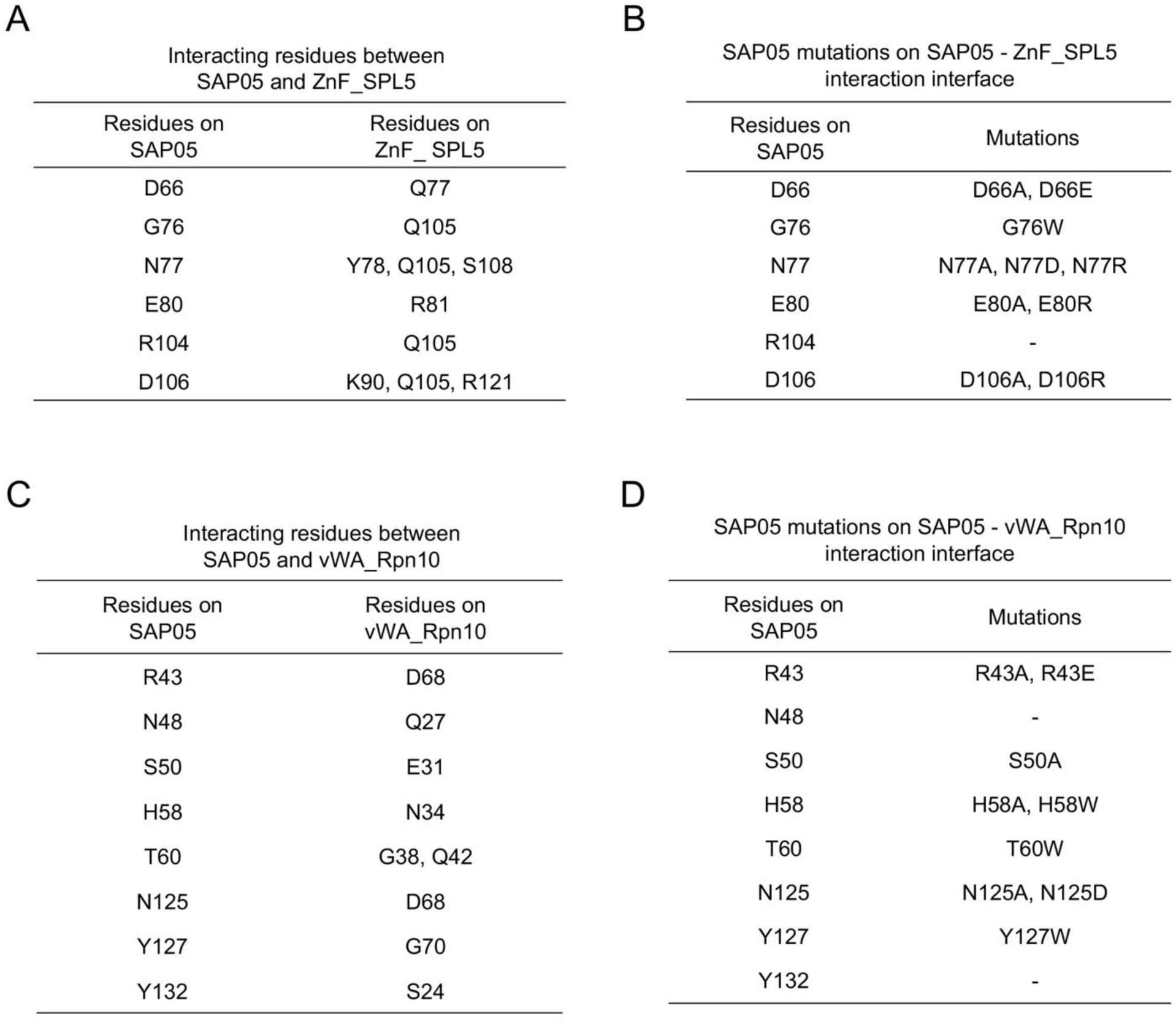
Residues that interact in the interfaces of SAP05 – ZnF_SPL5 (A) and SAP05 – vWA_Rpn10 (C) complexes and introduced mutations in SAP05 (B, D).

**Supplementary Fig. 3.**
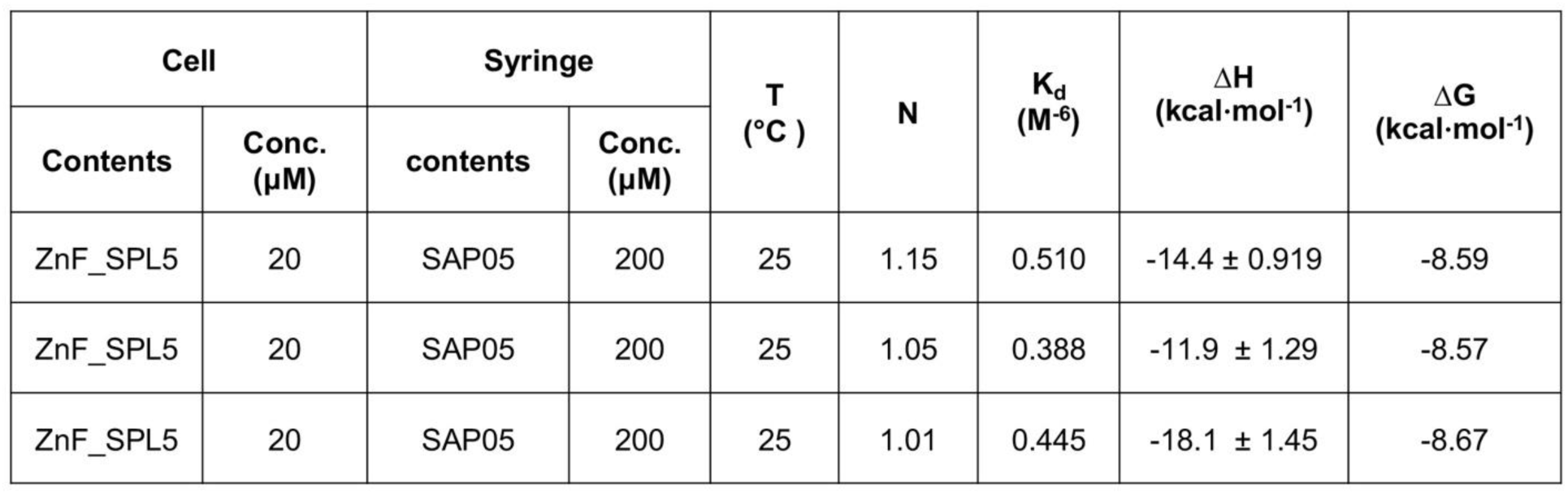
Thermodynamic parameters obtained from ITC binding tests for SAP05 and ZnF_SPL5 proteins. Three repeats of ITC tests by titrating SAP05 to ZnF_SPL5. Results of the first repeat is shown in Fig. 2C. Conc, concentration.

**Supplementary Fig. 4.**
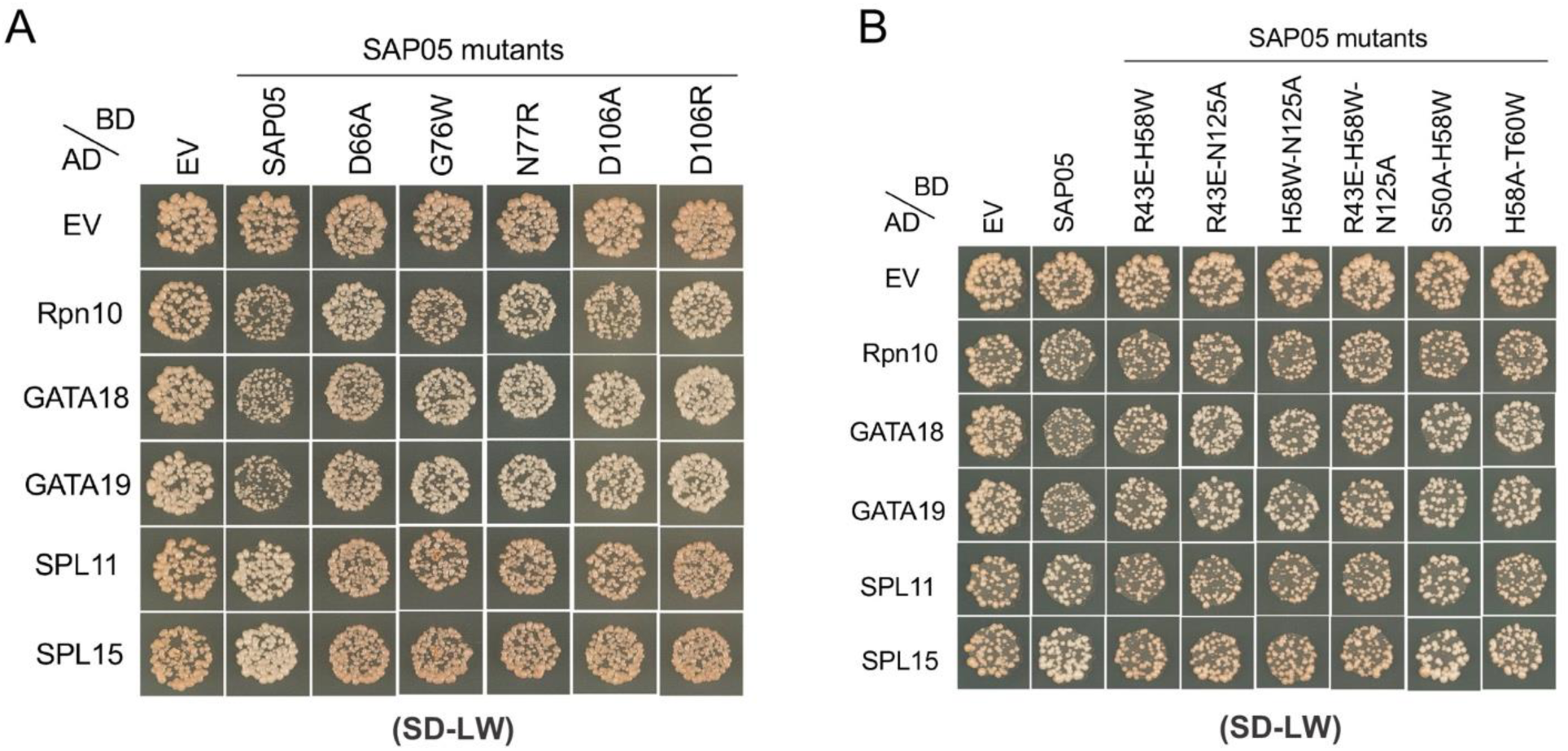
Yeast transformation controls for Y2H assays in the study, related to Figures 2 and 3. (A) Yeast growth on double dropout medium, SD-LW, lacking leucine and tryptophan indicating the expression of AD and BD constructs in Y2H assays for Figure 2E (A) and Figure 3C (B) in the main text.

**Supplementary Fig. 5.**
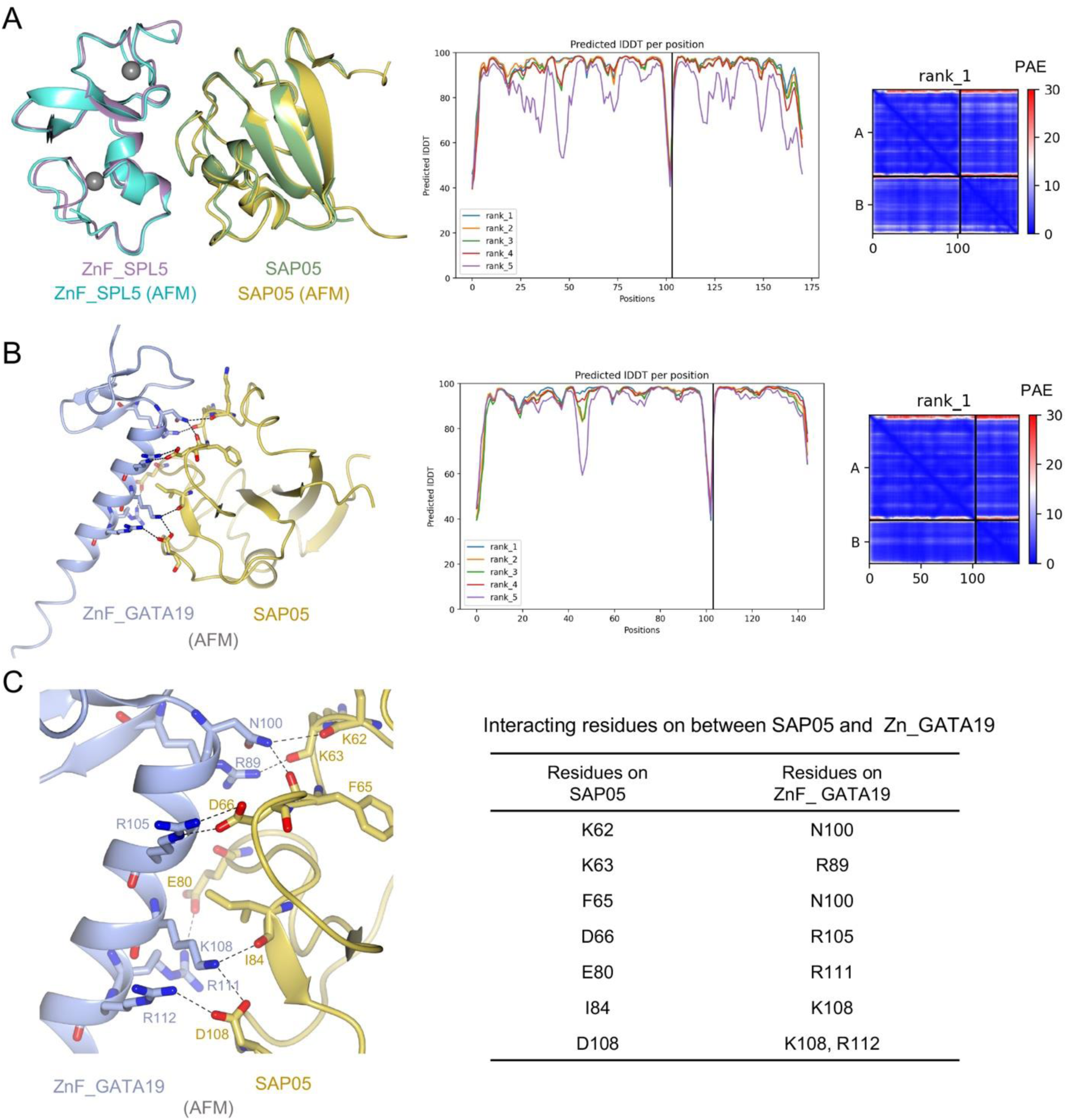
Structural prediction of SAP05 – ZnF_SPL5 and SAP05 – ZnF_GATA19 complexes. (A) Left, AlphaFold model (AFM) for SAP05 – ZnF_SPL5 complex (yellow – cyan) superimposed on the crystal structure of SAP05 – ZnF_SPL5 complex (green – purple). Middle, the predicted local Distance Difference Test (lDDT) value of SAP05 – ZnF_SPL5 model, showing the quality of predicted models by evaluating local distance differences to the reference. Best model (rank_1) was used for superimposition. Right, predicted aligned error (PAE) of the rank_1 model, showing the estimate of position error between predicted and true structures. Blue means lower error scores, red means higher error scores. (B) Left, AFM model (rank_1) of SAP05 – ZnF_GATA19 complex (yellow – light blue). Middle, predicted lDDT value of SAP05 – ZnF_GATA19 complex. Right, predicted aligned error (PAE) of the rank_1 model. (C) Left, enlarged view for predicted SAP05 – ZnF_GATA19 interaction interface. SAP05 residues and ZnF_GATA19 residues that involved for interaction with ZnF_GATA19 were labelled in yellow and light blue, respectively. Dashed line shows predicted interactions between SAP05 and ZnF_GATA19 from AFM model. Right, interacting residues between SAP05 and ZnF_GATA19 of predicted structure.

**Supplementary Fig. 6.**
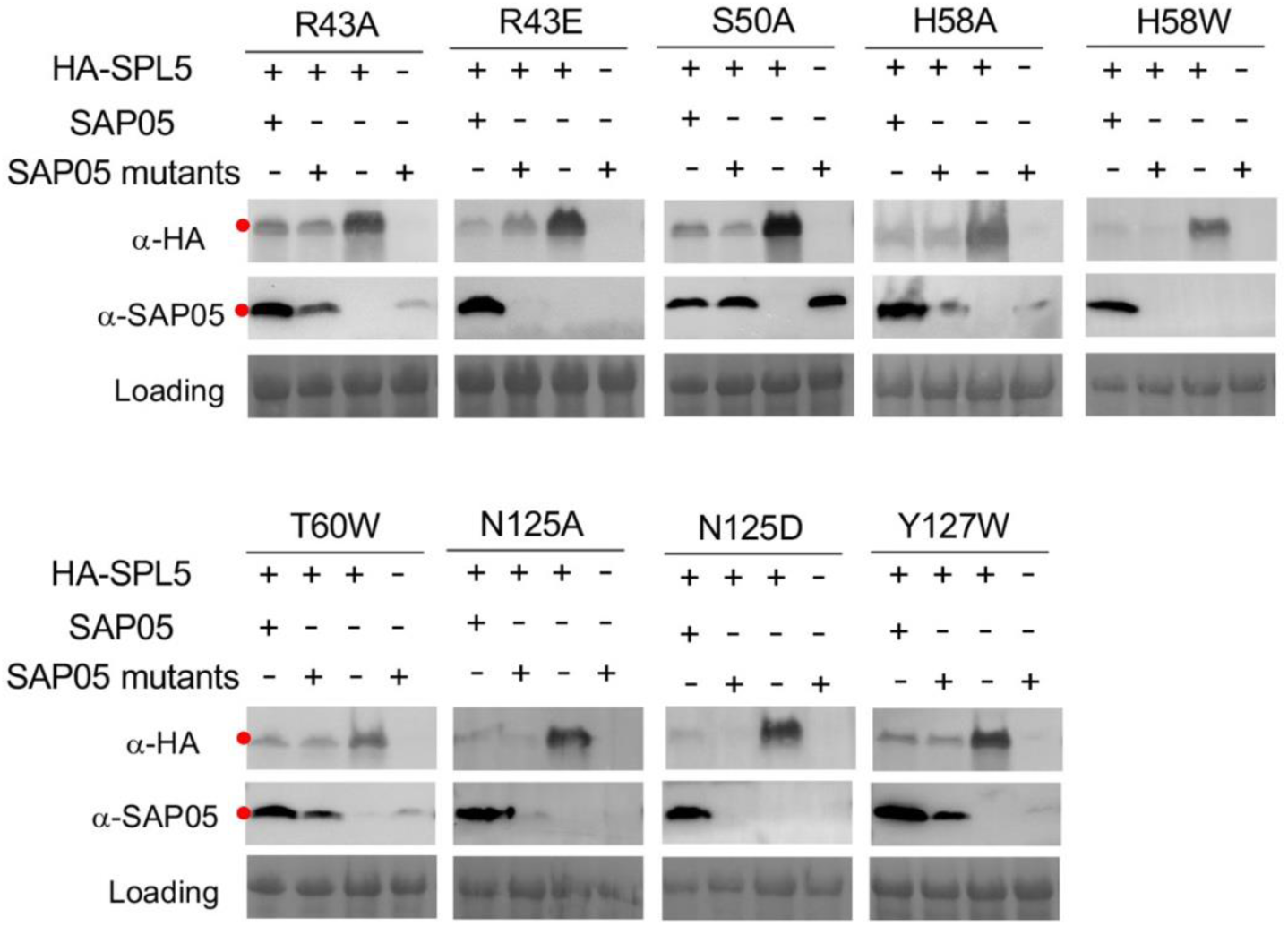
Western blot analysis of proteasomal degradation of SPL5 in presence of SAP05 wild-type or single mutants on sheet surface. Red dots on the blots indicate the expected sizes of the transiently expressed proteins in *N. benthamiana* leaves. Protein loading was visualized using Ponceau S staining. HA, Hemagglutinin.

**Supplementary Fig. 7.**
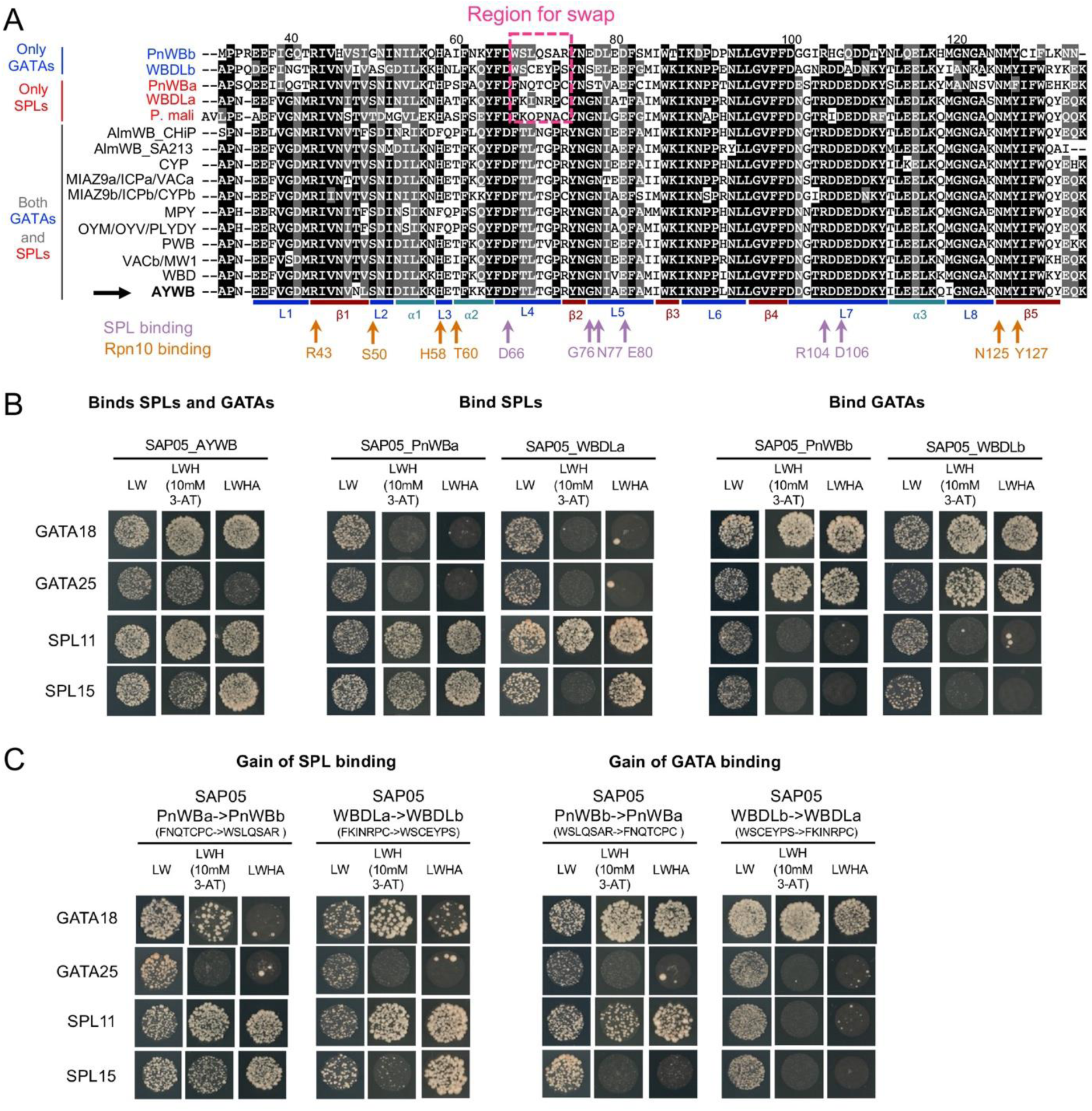
Conservation of SPL, GATA and Rpn10-interacting residues in SAP05 homologs. (A) Multiple sequence alignment of SAP05 homologs from different phytoplasma species showing that the Rpn10-binding or SPL-binding residues are conserved. Identical and conserved amino acid residues are denoted by black and gray backgrounds, respectively. Homologs binding only GATAs, only SPLs, and both GATAs and SPLs are displayed in blue, red and black, respectively, at the left of the alignment. The secondary structure of AYWB SAP05 is highlighted at the bottom (color-matched with Fig. 1A-B). SAP05 residues involved in SPL binding (purple) or Rpn10 binding (orange) are pointed with arrows (color-matched with Fig. 1C-F). The region used for swap in (C) is marked with a pink dashed rectangle. (B) Y2H assay showing SAP05 homologs that specifically bind SPLs or GATAs. (C) Y2H assay showing that swapping loop 4 sequences from GATA-binding SAP05 homologs contributes to SPL binding, and swapping loop 4 sequences from SPL-binding SAP05 homologs to GATA binding. (B, C) Yeasts were grown on drop-out media lacking L, W, H or A, or in the presence of 3-amino-1,2,4-triazole (3-AT).

**Supplementary Fig. 8.**
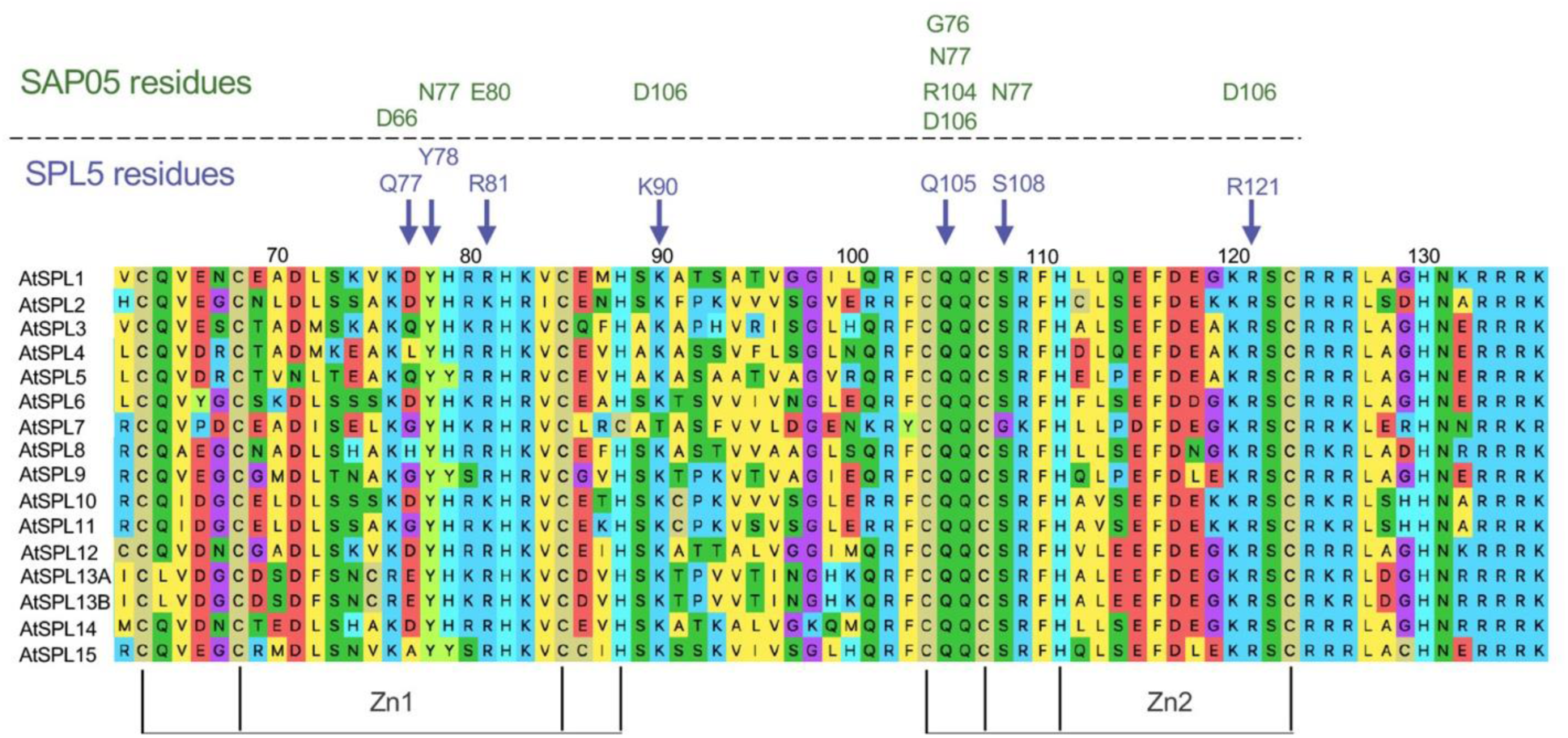
Conservation level of *A. thaliana* SPL5 residues that interact with SAP05 among *A. thaliana* SPL family members. The multiple sequence alignment was generated with MUSCLE. The conserved SPL residues involved in the interaction with SAP05 are indicated with purple arrows and the corresponding SAP05 residues are shown in green. Accession numbers for AtSPLs shown are: SPL1, AT2G47070; SPL2, AT5G43270; SPL3, AT2G33810; SPL4, AT1G53160; SPL5, AT3G15270; SPL6, AT1G69170; SPL7, AT5G18830; SPL8, AT1G02065; SPL9, AT2G42200; SPL10, AT1G27370; SPL11, AT1G27360; SPL12, AT3G60030; SPL13A, AT5G50570; SPL13B, AT5G50670; SPL14, AT1G20980; SPL15, AT3G57920.

**Supplementary Fig. 9.**
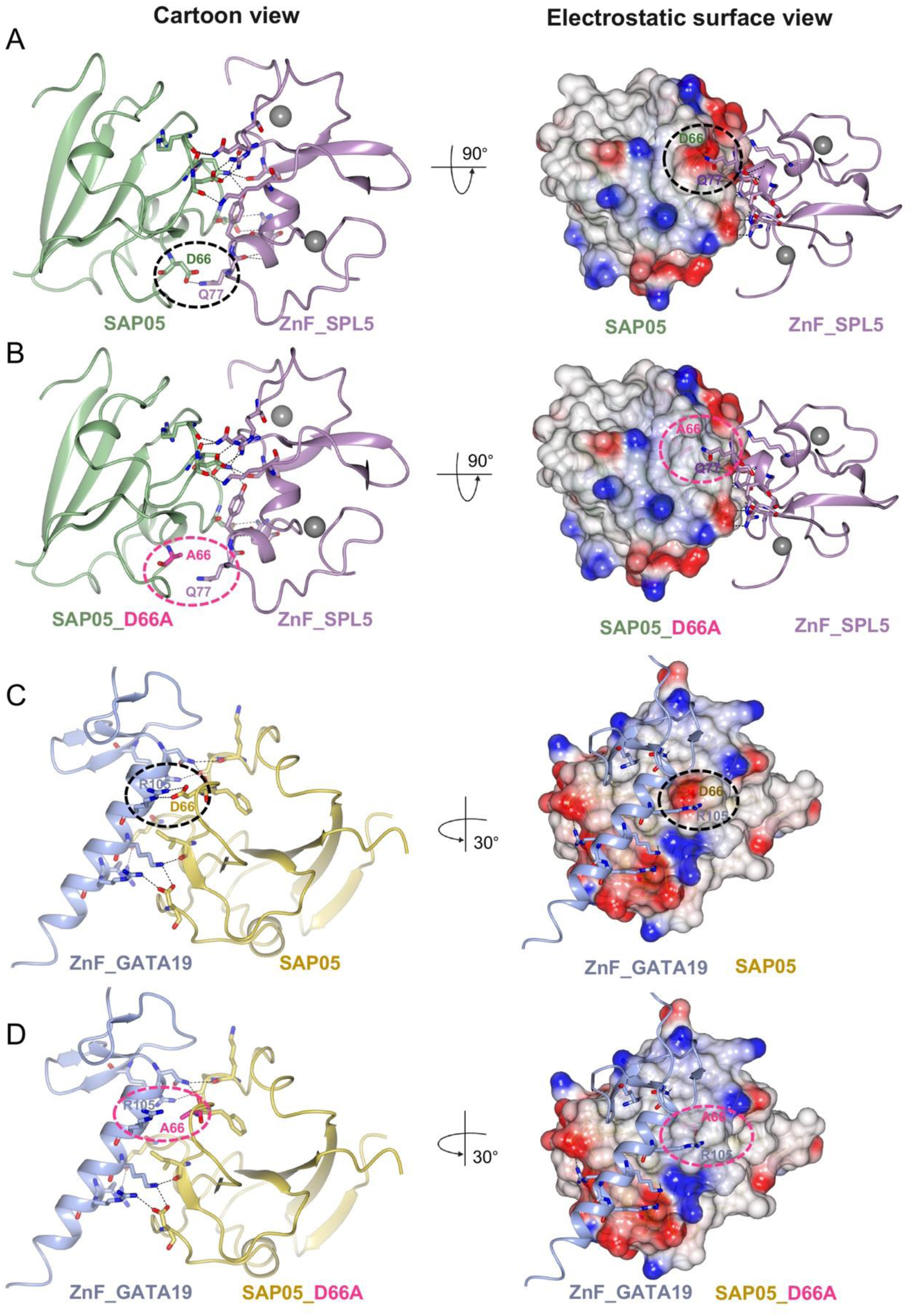
SAP05 D66 creates an electronegative surface at the binding interface with zinc-fingers of SPL and GATA. (A) ZnF_SPL5 Q77 interacts with SAP05 D66 (left), which creates an electronegative surface at the binding interface with SPL5 (right). (B) SPL5 Q77 doesn’t interact with SAP05 D66A (left) and the electronegativity of the interaction surface is reduced (right). (C) ZnF_GATA19 R105 interacts with SAP05 D66 (left) based on the AlphaFold-predicted top-ranked complex structure, and D66 creates an electronegative surface at the binding interface with GATA19 (right). (D) GATA19 R105 isn’t predicted to interact with SAP05 D66A (left) and the electronegativity of the interaction surface is reduced (right).

**Supplementary Fig. 10.**
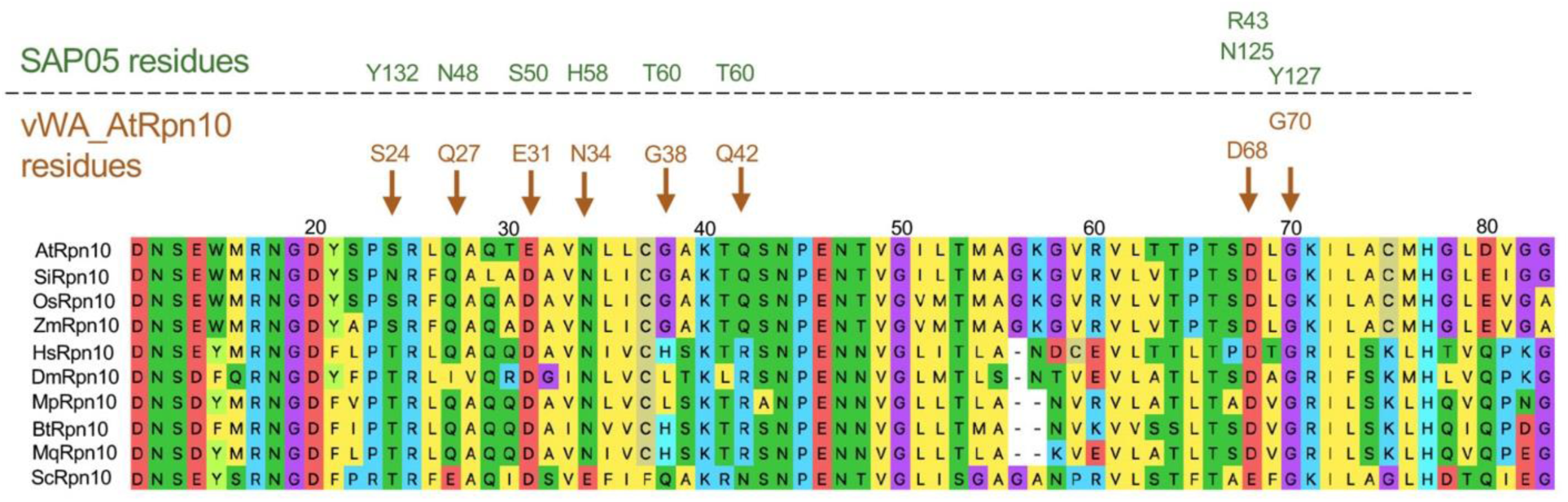
Conservation level of *A. thaliana* Rpn10 residues that interact with SAP05 among Rpn10 homologs from other organisms. The multiple sequence alignment was generated with MUSCLE. The conserved vWA_Rpn10 residues involved in interaction with SAP05 are indicated with orange arrows, and the corresponding SAP05 residues are shown on top in green. Sequence alignment was conducted with MUSCLE in MEGA11 software. AtRpn10, *Arabidopsis thaliana* Rpn10 (Uniprot ID: P55034); SlRpn10, *Solanum lycopersicum* Rpn10 (Uniprot ID: A0A3Q7F6N7); OsRpn10, *Oryza sativa* Rpn10 (Uniprot ID: O82143); ZmRpn10, *Zea mays* Rpn10 (Uniprot ID: B6TK61); HsRpn10, *Homo sapiens* Rpn10 (Uniprot ID: Q5VWC4); DmRpn10, *Drosophila melanogaster* Rpn10 (Uniprot ID: P55035); MpRpn10, *Myzus persicae* Rpn 10 (GenBank: XP_022181722.1); BtRpn10, *Bemisia tabaci* Rpn10 (GenBank: XP_018915695); MqRpn10, *Macrosteles quadrilineatus* Rpn10; ScRpn10, *Saccharomyces cerevisiae* Rpn10 (Uniprot ID: P38886).

**Supplementary Fig. 11.**
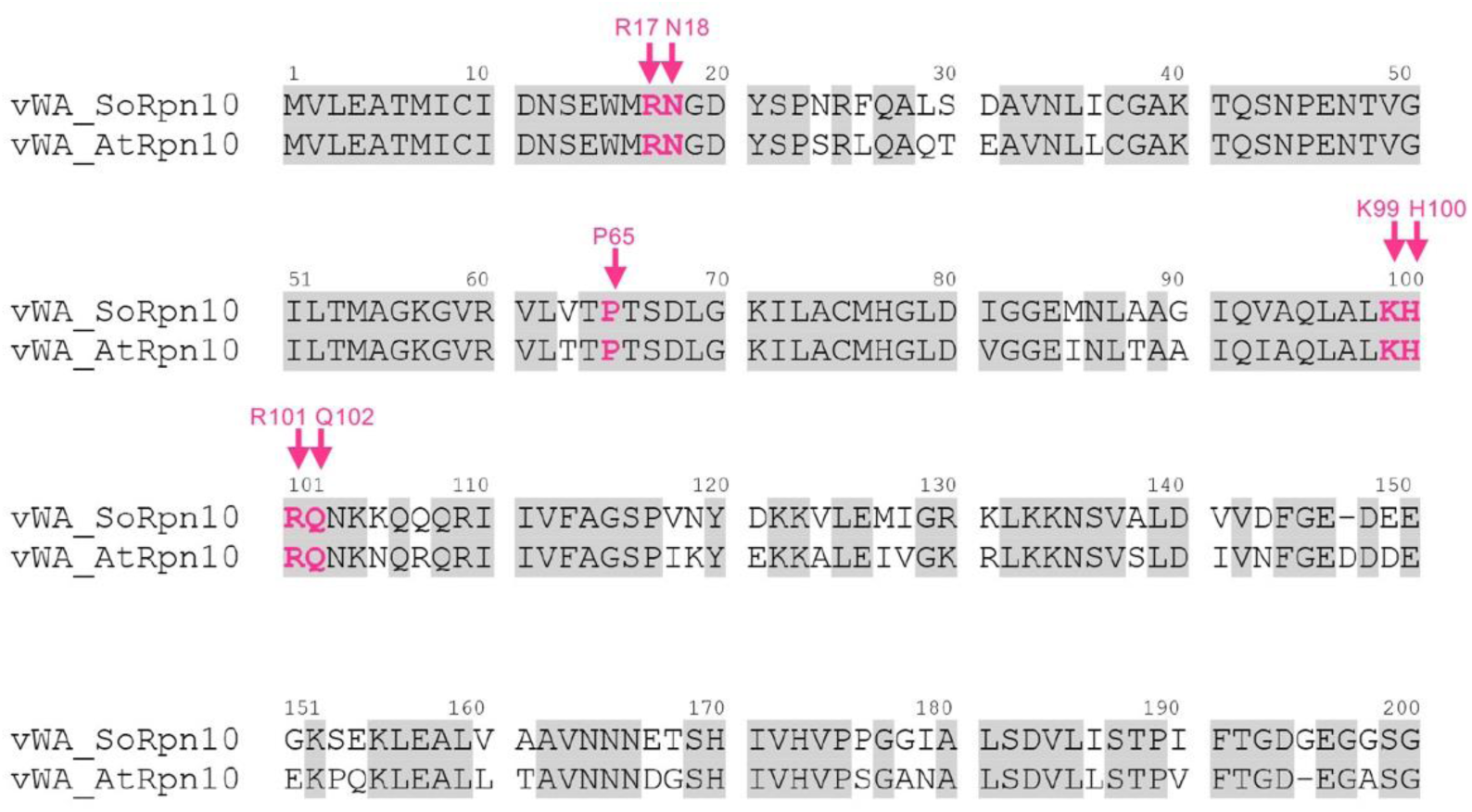
Sequence alignment of vWA_Rpn10 from *Spinacia oleracea* and *A. thaliana*. The conserved vWA_Rpn10 residues are marked with grey background. Residues interacting with 26S proteasome are marked with pink arrows. The vWA_SoRpn10 sequence was extracted from the structure of spinach 19S proteasome (PDB 8AMZ). Sequence alignment was conducted with MUSCLE in MEGA11 software and residues interacting with 26S proteasome were analyzed with CCP4mg.

**Supplementary Fig. 12.**
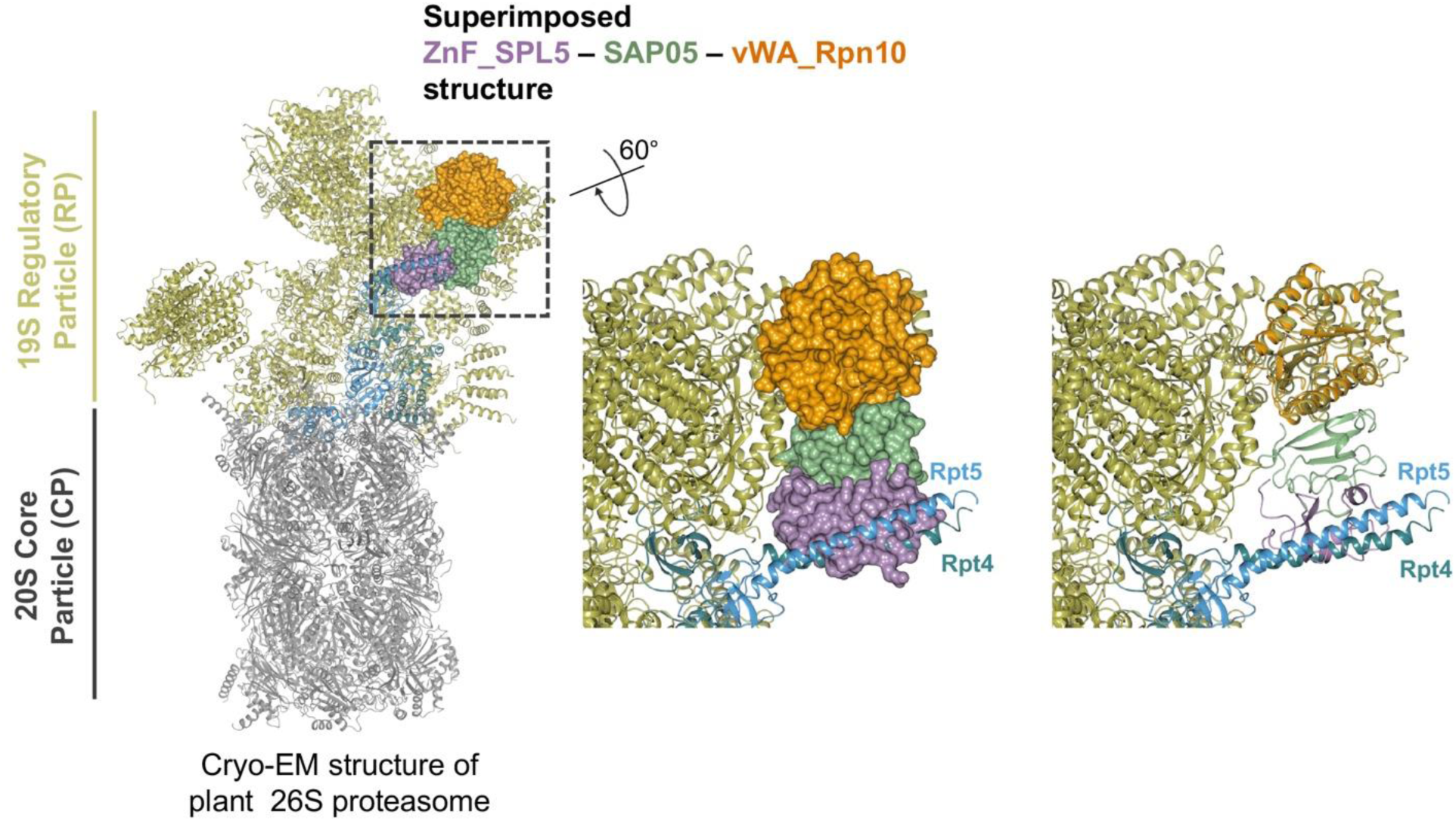
SAP05 – ZnF_SPL5 binding to vWA_Rpn10 clashes with plant 26S proteasome. Structural superimposition of the ternary structure of ZnF_SPL5 – SAP05 – vWA_Rpn10 on the spinach 26S proteasome (PDB 7QVE and PDB 8AMZ) shows steric clashes (left). A zoom-in view of the superimposed structures highlights the clashes between ZnF_SPL5 and two ⍺-helices of 19S RP, which are homologs of *A. thaliana* Rpt4 (turquoise) and Rpt5 (blue). The ZnF_SPL5 – SAP05 – vWA_Rpn10 complex is depicted as surface (middle) and cartoon (right) presentations.

## References

Bard, J.A.M., Goodall, E.A., Greene, E.R., Jonsson, E., Dong, K.C., and Martin, A. (2018). Structure and Function of the 26S Proteasome. Annu. Rev. Biochem. 87, 697–724.

Barrett, A.J. (2001). Proteases. Curr. Protoc. Protein Sci. Chapter 21, Unit 21.1. 10.1002/0471140864.ps2101s21.

Bastos, M., and Velazquez-Campoy, A. (2021). Isothermal titration calorimetry (ITC): a standard operating procedure (SOP). Eur. Biophys. J. 50, 363–371.

Baumeister, W., Walz, J., Zühl, F., and Seemüller, E. (1998). The proteasome: paradigm of a self-compartmentalizing protease. Cell 92, 367–380.

Benoit, R.M., Wilhelm, R.N., Scherer-Becker, D., and Ostermeier, C. (2006). An improved method for fast, robust, and seamless integration of DNA fragments into multiple plasmids. Protein Expr. Purif. 45, 66–71.

Berrow, N.S., Alderton, D., Sainsbury, S., Nettleship, J., Assenberg, R., Rahman, N., Stuart, D.I., and Owens, R.J. (2007). A versatile ligation-independent cloning method suitable for high-throughput expression screening applications. Nucleic Acids Res. 35, e45.

Bhattacharyya, S., Yu, H., Mim, C., and Matouschek, A. (2014). Regulated protein turnover: snapshots of the proteasome in action. Nat. Rev. Mol. Cell Biol. 15, 122–133.

Bird, L.E., Rada, H., Flanagan, J., Diprose, J.M., Gilbert, R.J., and Owens, R.J. (2014). Application of In-Fusion™ cloning for the parallel construction of E. coli expression vectors. Methods Mol. Biol. 1116, 209–234.

Bogdanove, A.J., and Voytas, D.F. (2011). TAL effectors: customizable proteins for DNA targeting. Science (New York, NY) 333, 1843–1846.

Buneeva, O.A., and Medvedev, A.E. (2018). Ubiquitin-independent protein degradation in proteasomes. Biomed. Khim. 64, 134–148.

Chen, L., Shinde, U., Ortolan, T.G., and Madura, K. (2001). Ubiquitin-associated (UBA) domains in Rad23 bind ubiquitin and promote inhibition of multi-ubiquitin chain assembly. EMBO Rep. 2(10):933–8.

Chen, X., Dorris, Z., Shi, D., Huang, R.K., Khant, H., Fox, T., de Val, N., Williams, D., Zhang, P., and Walters, K.J. (2020). Cryo-EM Reveals Unanchored M1-Ubiquitin Chain Binding at hRpn11 of the 26S Proteasome. Structure 28(11):1206–1217.e4.

Ciechanover, A. (2017). Intracellular protein degradation: From a vague idea thru the lysosome and the ubiquitin-proteasome system and onto human diseases and drug targeting. Best Pract. Res. Clin. Haematol. 30, 341–355.

Collins, G.A., and Goldberg, A.L. (2017). The Logic of the 26S Proteasome. Cell 169, 792–806.

Darwin, K.H. (2009). Prokaryotic ubiquitin-like protein (Pup), proteasomes and pathogenesis. Nat. Rev. Microbiol. 7, 485–491. 10.1038/nrmicro2148.

Davis, C., Spaller, B.L., and Matouschek, A. (2021). Mechanisms of substrate recognition by the 26S proteasome. Curr. Opin. Struct. Biol. 67, 161–169.

Dereeper, A., Guignon, V., Blanc, G., Audic, S., Buffet, S., Chevenet, F., Dufayard, J.F., Guindon, S., Lefort, V., Lescot, M., Claverie, J.M., and Gascuel, O. (2008). Phylogeny.fr: robust phylogenetic analysis for the non-specialist. Nucleic Acids Res. 36: W465–9.

Du, T., Song, Y., Ray, A., Wan, X., Yao, Y., Samur, M.K., Shen, C., Penailillo, J., Sewastianik, T., Tai, Y.T., Fulciniti, M., Munshi, N.C., Wu, H., Carrasco, R.D., Chauhan, D., and Anderson, K.C. (2023). Ubiquitin receptor PSMD4/Rpn10 is a novel therapeutic target in multiple myeloma. Blood. 141(21):2599–2614.

Elsasser, S., Chandler-Militello, D., Müller, B., Hanna, J., Finley, D. (2004). Rad23 and Rpn10 serve as alternative ubiquitin receptors for the proteasome. J. Biol. Chem. 279(26):26817–22.

Emsley, P., Lohkamp, B., Scott, W.G., and Cowtan, K. (2010). Features and development of Coot. Acta Crystallogr. D Biol. Crystallogr. 66, 486–501.

Evans, P.R., and Murshudov, G.N. (2013). How good are my data and what is the resolution? Acta Crystallogr. D Biol. Crystallogr. 69, 1204–1214.

Fatimababy, A.S., Lin, Y.L., Usharani, R., Radjacommare, R., Wang, H.T., Tsai, H.L., Lee, Y., and Fu, H. (2010). Cross-species divergence of the major recognition pathways of ubiquitylated substrates for ubiquitin/26S proteasome-mediated proteolysis. FEBS J. 277(3):796–816.

Finley, D. (2009). Recognition and processing of ubiquitin-protein conjugates by the proteasome. Annu. Rev. Biochem. 78, 477–513.

Glickman, M.H., Rubin, D.M., Coux, O., Wefes, I., Pfeifer, G., Cjeka, Z., Baumeister, W., Fried, V.A., and Finley, D., (1998). A subcomplex of the proteasome regulatory particle required for ubiquitin-conjugate degradation and related to the COP9-signalosome and eIF3. Cell 94(5), pp.615–623.

Hamazaki, J., Sasaki, K., Kawahara, H., Hisanaga, S., Tanaka, K., and Murata, S. (2007). Rpn10-mediated degradation of ubiquitinated proteins is essential for mouse development. Mol. Cell Biol. 27(19):6629–38.

Hartley, J.L., Temple, G.F., and Brasch, M.A. (2000). DNA cloning using in vitro site-specific recombination. Genome Res. 10, 1788–1795.

Hogenhout, S.A., Van der Hoorn, R.A., Terauchi, R., and Kamoun, S. (2009). Emerging concepts in effector biology of plant-associated organisms. Mol. Plant Microbe Interact. 22, 115–122.

Huang, W., MacLean, A.M., Sugio, A., Maqbool, A., Busscher, M., Cho, S.T., Kamoun, S., Kuo, C.H., Immink, R.G.H., and Hogenhout, S.A. (2021). Parasitic modulation of host development by ubiquitin-independent protein degradation. Cell 184(20):5201–5214.e12.

Huang, W., and Hogenhout, S.A. (2022). Interfering with plant developmental timing promotes susceptibility to insect vectors of a bacterial parasite. Biorxiv, https://doi.org/10.1101/2022.03.30.486463.

Inobe, T., and Matouschek, A. (2014). Paradigms of protein degradation by the proteasome. Curr. Opin. Struct. Biol. 24, 156–164.

Jumper, J., Evans, R., Pritzel, A., Green, T., Figurnov, M., Ronneberger, O., Tunyasuvunakool, K., Bates, R., Žídek, A., Potapenko, A., et al. (2021). Highly accurate protein structure prediction with AlphaFold. Nature 596, 583–589.

Kandolf, S., Grishkovskaya, I., Belačić, K., Bolhuis, D.L., Amann, S., Foster, B., Imre, R., Mechtler, K., Schleiffer, A., Tagare, H.D., et al. (2022). Cryo-EM structure of the plant 26S proteasome. Plant Commun. 3, 100310.

Kitazawa, Y., Iwabuchi, N., Maejima, K., Sasano, M., Matsumoto, O., Koinuma, H., Tokuda, R., Suzuki, M., Oshima, K., Namba, S., and Yamaji, Y. (2022). A phytoplasma effector acts as a ubiquitin-like mediator between floral MADS-box proteins and proteasome shuttle proteins. Plant Cell 34, 1709–1723.

Komander, D., and Rape, M. (2012). The ubiquitin code. Annu. Rev. Biochem. 81, 203–229.

Langin, G., and Üstün, S. (2023). A Pipeline to Monitor Proteasome Homeostasis in Plants. Methods Mol. Biol. 2581, 351–363.

Leestemaker, Y., and Ovaa, H. (2017). Tools to investigate the ubiquitin proteasome system. Drug Discov. Today Technol. 26, 25–31.

Liang, X., Peng, L., Baek, C.H., and Katzen, F. (2013). Single step BP/LR combined Gateway reactions. BioTechniques 55, 265–268.

Lu, X., Sabbasani, V.R., Osei-Amponsa, V., Evans, C.N., King, J.C., Tarasov, S.G., Dyba, M., Das, S., Chan, K.C., Schwieters, C.D., Choudhari, S., Fromont, C., Zhao, Y., Tran, B., Chen, X., Matsuo, H., Andresson, T., Chari, R., Swenson, R.E., Tarasova, N.I., and Walters, K.J. (2021). Structure-guided bifunctional molecules hit a DEUBAD-lacking hRpn13 species upregulated in multiple myeloma. Nat. Commun. 12(1):7318.

MacLean, A.M., Orlovskis, Z., Kowitwanich, K., Zdziarska, A.M., Angenent, G.C., Immink, R.G., and Hogenhout, S.A. (2014). Phytoplasma effector SAP54 hijacks plant reproduction by degrading MADS-box proteins and promotes insect colonization in a RAD23-dependent manner. PLoS Biol. 12, e1001835.

Mao, Y. (2021). Structure, Dynamics and Function of the 26S Proteasome. Subcell. Biochem. 96, 1–151.

Matyskiela, M.E., Lander, G.C., and Martin, A. (2013). Conformational switching of the 26S proteasome enables substrate degradation. Nat. Struct. Mol. Biol. 20(7):781–788.

McCoy, A.J., Grosse-Kunstleve, R.W., Adams, P.D., Winn, M.D., Storoni, L.C., and Read, R.J. (2007). Phaser crystallographic software. J. Appl. Crystallogr. 40, 658–674.

McNicholas S., Potterton E., Wilson K.S. and Noble M.E. (2011). Presenting your structures: the CCP4mg molecular-graphics software. Acta Crystallogr. D Biol. Crystallogr. 67, 386–394.

Mojica, F.J., and Rodriguez-Valera, F. (2016). The discovery of CRISPR in archaea and bacteria. FEBS J. 283, 3162–3169.

Müller, A.U., and Weber-Ban, E. (2019). The Bacterial Proteasome at the Core of Diverse Degradation Pathways. Front Mol. Biosci. 6, 23.

Murata, S., Yashiroda, H., and Tanaka, K. (2009). Molecular mechanisms of proteasome assembly. Nat. Rev. Mol. Cell Biol. 10, 104–115.

Murshudov, G.N., Skubak, P., Lebedev, A.A., Pannu, N.S., Steiner, R.A., Nicholls, R.A., Winn, M.D., Long, F., and Vagin, A.A. (2011). REFMAC5 for the refinement of macromolecular crystal structures. Acta Crystallogr. D Biol. Crystallogr. 67, 355–367.

Nelson, M.D., and Fitch, D.H. (2011). Overlap extension PCR: an efficient method for transgene construction. Methods Mol. Biol. 772, 459–470.

Rani, N., Aichem, A., Schmidtke, G., Kreft, S.G., and Groettrup, M. (2012). FAT10 and NUB1L bind to the VWA domain of Rpn10 and Rpn1 to enable proteasome-mediated proteolysis. Nat. Commun. 3:749.

Richard, E., Michael, O.N., Alexander, P., Natasha, A., Andrew, S., Tim, G., Augustin, Ž., Russ, B., Sam, B., Jason, Y., et al. (2022). Protein complex prediction with AlphaFold-Multimer. bioRxiv, 2021.2010.2004.463034.

Rousseau, A., and Bertolotti, A. (2018). Regulation of proteasome assembly and activity in health and disease. Nat. Rev. Mol. Cell Biol. 19, 697–712.

Saeki, Y. (2017). Ubiquitin recognition by the proteasome. J. Biochem. 161, 113–124.

Sakata, E., Bohn, S., Mihalache, O., Kiss, P., Beck, F., Nagy, I., Nickell, S., Tanaka, K., Saeki, Y., Förster, F., and Baumeister, W. (2012). Localization of the proteasomal ubiquitin receptors Rpn10 and Rpn13 by electron cryomicroscopy. Proc. Natl. Acad. Sci. U S A. 109(5):1479-84.

Skubak, P., Arac, D., Bowler, M.W., Correia, A.R., Hoelz, A., Larsen, S., Leonard, G.A., McCarthy, A.A., McSweeney, S., Mueller-Dieckmann C., Otten, H., Salzman, G., and Pannu, N.S. (2018). A new MR-SAD algorithm for the automatic building of protein models from low-resolution X-ray data and a poor starting model. IUCrJ 5, 166–171.

Snoberger, A., Brettrager, E.J., and Smith. D.M. (2018). Conformational switching in the coiled-coil domains of a proteasomal ATPase regulates substrate processing. Nat. Commun. 9(1):2374.

Tanaka, K. (2009). The proteasome: overview of structure and functions. Proc. Jpn. Acad. Ser. B Phys. Biol. Sci. 85, 12–36.

Verma, R., Oania, R., Graumann, J., Deshaies, R.J. (2004). Multiubiquitin chain receptors define a layer of substrate selectivity in the ubiquitin-proteasome system. Cell 118(1):99–110.

Wang, Q., Young, P., and Walters, K. J. (2005). Structure of S5a bound to monoubiquitin provides a model for polyubiquitin recognition. J. Mol. Biol. 348(3):727–39.

Waterhouse, A., Bertoni, M., Bienert, S., Studer, G., Tauriello, G., Gumienny, R., Heer, F.T., de Beer, T.A.P., Rempfer, C., Bordoli, L., Lepore, R., and Schwede, T. (2018). SWISS-MODEL: homology modelling of protein structures and complexes. Nucleic Acids Res. 46, W296–W303.

Wei, P., Wong, W.W., Park, J.S., Corcoran, E.E., Peisajovich, S.G., Onuffer, J.J., Weiss, A., and Lim, W.A. (2012). Bacterial virulence proteins as tools to rewire kinase pathways in yeast and immune cells. Nature 488, 384–388.

Wickham, H. (2011). Ggplot2. Wiley Interdisciplinary Reviews: Computational Statistics. 3, 2, pp. 180––185.

Winn, M.D., et al. (2011). Overview of the CCP4 suite and current developments. Acta Crystallogr. D Biol. Crystallogr. 67, 235–242.

Winter, G. (2010) Xia2: an expert system for macromolecular crystallography data reduction. J. Appl. Crystallogr. 43, 186–190.

Winter, G., Waterman, D.G., Parkhurst, J.M., Brewster, A.S., Gildea, R.J., Gerstel, M., Fuentes-Montero, L., Vollmar, M., Michels-Clark, T., Young, I.D., Sauter, N.K., and Evans, G. (2018). DIALS: implementation and evaluation of a new integration package. Acta Crystallogr. D Struct. Biol. 74(2):85–97.

Xu, F.Q., and Xue, H.W. (2019). The ubiquitin-proteasome system in plant responses to environments. Plant Cell Environ. 42, 2931–2944.

Yoo, S.D., Cho, Y.H., and Sheen, J. (2007). Arabidopsis mesophyll protoplasts: a versatile cell system for transient gene expression analysis. Nat. Protoc. 2(7): 1565–72.

Yu, H., and Matouschek, A. (2017). Recognition of Client Proteins by the Proteasome. Annu. Rev. Biophys. 46, 149–173.

Zambelli, B. (2019). Characterization of Enzymatic Reactions Using ITC. Methods Mol. Biol. 1964, 251–266.

Zhu, Y., Wang, W.L., Yu, D., Ouyang, Q., Lu, Y., and Mao, Y. (2018). Structural mechanism for nucleotide-driven remodeling of the AAA-ATPase unfoldase in the activated human 26S proteasome. Nat. Commun. 9, 1360.

